# The role of chromosomal inversions in local adaptation and ecotype formation across a variable landscape in an intertidal snail

**DOI:** 10.64898/2026.10.19.760124

**Authors:** James Reeve, Olga Ortega-Martinez, Per Bergström, Eva Koch, Kerstin Johannesson, Roger K. Butlin

## Abstract

Adaption to environmental variation can be discrete or continuous, responding to sharp or gradual habitat transitions. Chromosomal inversions accentuate these adaptive responses as multiple loci respond in tandem across a habitat transition. Earlier studies of *Littorina saxatilis* (Olivi, 1792) have shown that three ecotypes segregate in different habitats supported by polymorphisms in a few inversions. However, many more large inversions are polymorphic in this species and potentially have adaptive effects in response to uncharacterised environmental gradients. We sampled 200 random shores representing all kinds of shore habitats present in a 12x12 km area of a Swedish archipelago, with *L. saxatilis* found at 196 sites. Environment and shell morphology were summarised using dimension reduction analyses, and Bayesian regression models were used to analyse the relationships between phenotype, environment and inversions. Phenotypic variation was a continuum with previously described ecotypes positioned in different parts. 18 inversions responded to at least one dimension of environmental variation, and over (12) half influenced the measured phenotypes. Overall, inversion arrangements explained 15% of phenotypic variation, with limited direct contributions from the environment (<1%). Chromosomal inversions thus are a major factor for local adaptation in *L. saxatilis*, supporting adaptation across a landscape more complex than the previously described ecotypes.

## Introduction

Local adaptation occurs within a complex landscape of environmental variation, yet it is typically studied in the context of simple gradients or contrasting habitat classes. Spatial variation in adaptive phenotypes and genotypes depends on the interaction between habitat-dependent selection pressures and the homogenising effects of gene flow. A commonly studied pattern is broad-scale clinal variation associated with environmental gradients that are wide relative to dispersal (‘ecoclines’). Alternatively, distinct phenotypes (“ecotypes”) associated with discrete habitat types and sharp genetic clines, may form at habitat boundaries where multiple environmental changes occur together (“ecotones”; Turesson 1922; Savolainen, et al. 2013; Johannesson et al., 2026). Clines in many traits and genes may be coupled at ecotones (Bierne et al., 2011), which can become a problem when trying to identify adaptive loci. While ecotypes offer strong evidence for local adaptation, separating out independent adaptive responses among ecotype pairs can be difficult unless coupling across an ecotone can be broken.

Continuous and discrete environmental patterns can occur together where the frequencies of different local patch types vary on a larger spatial scale, producing steep changes in traits and genotype frequencies at local boundaries but a shallower cline when viewed at low resolution (Harrison & Rand, 1989; Ross & Harrison, 2002). For example, a smooth ecocline running across Connecticut (USA) connects two species of field cricket, but at a finer scale these species are demarcated by ecotones between distinct soil types (Ross & Harrison, 2002). This smoothing out of patterns occurs as the local genetic structure and habitat conditions get blurred into population level trends at broader scales (Levin, 1992; Rogivue et al., 2025). The resolution of populations across gradients of environmental variation depends partly on the dispersal distance of the focal species, which determines the perceived grain of the environment and the scale of local adaptation (Levin 1976; Slatkin 1973). In addition, the environmental drivers of local adaptation identified by genotype-environment associations also change with sampling scale (Guillaume et al., 2024; Guillaume, 2024). Our conclusions about the nature of local adaptation heavily depend on how we sample variable landscapes (Lotterhos & Whitlock, 2015). Focused sampling that compares contrasting habitats tends to identify discrete populations, while random sampling provides a broader perspective.

Ecotypes are often thought of as symptoms of strong local adaptation. They have been studied to determine the genetic architecture associated with local adaptation, as a presumed early step in speciation (Lucek, 2012; Lowry et al., 2014; Raszick & Song, 2016; Stronen et al., 2022). Typically, ecotype studies focus their sampling efforts on patches of distinct habitat (e.g., *Helianthus petiolaris* [Todesco et al., 2020], and *Timema* stick insects [Comeault et al., 2015]) or the extreme ends of environmental gradients (e.g., threespine stickleback [Jones et al., 2012; Ravinet et al., 2015], switchgrass [Lowry et al., 2014], and deer mice [Hager et al., 2022]). Discrete ecotypes are most likely to evolve where the environmental variation is patchy, especially where multiple environmental factors vary together leading to differentiation of multiple phenotypic traits or genetic markers (Johannesson et al. 2026). The apparent discreteness of ecotype boundaries may also depend on sampling design.

Chromosomal inversions facilitate divergence among populations connected by gene flow by maintaining linkage disequilibrium among locally-adapted alleles (Kirkpatrick & Barton, 2006). There are multiple examples where inversion arrangement frequencies differ between ecotypes, such as Atlantic cod (Matschiner et al., 2022), European anchovy (Le Moan et al., 2016), threespine sticklebacks (Jones et al., 2012), *Timema* stick insects (Lucek et al., 2019), rough periwinkle snails (Faria et al., 2019), deer mice (Hager et al., 2022; Harringmeyer & Hoekstra, 2022), prairie sunflowers (Todesco et al., 2020), and monkey flowers (Coughlan & Willis, 2019; Kollar et al., 2023). In some cases, adaptive loci have been identified within these inversions (Kirubakaran et al., 2016; Lee et al., 2017; Coughlan & Willis, 2019; Kollar et al., 2023; Soudi et al., 2023). Due to strong linkage disequilibrium, the combined effects of all loci within an inversion can be treated as a single locus (Thompson & Jiggins, 2014), although genetic variation may accumulate within each arrangement (Berdan et al., 2023). This has a practical disadvantage as the loci contributing to local adaptation are hard to differentiate from linked neutral loci within an inversion, or from other non-adaptive loci which contribute to reproductive isolation. However, because of linkage between adaptive loci, inversions are often under strong selection in response to environmental gradients (Wellenreuther & Bernatchez, 2018; Westram et al., 2022; Berdan et al., 2023). Studying inversion arrangement frequencies across a single environmental transition will typically amplify the coupling of loci, as they will all become statistically associated with the major habitat transition (Bierne et al., 2011). Sampling across multiple environments in a landscape may help to decouple these genotype-phenotype-environment associations.

This project aimed to decouple the adaptive effects across a hybrid zone in the rough periwinkle (*Littorina saxatilis,* Olivi, 1792), by randomly sampling among all available habitats within a local geographic region. *Littorina saxatilis* has been repeatedly studied to understand the ecological and molecular mechanisms involved in local adaptation and ecotype formation (Johannesson et al., 2010; Faria et al. 2019; Morales et al., 2019; Reeve et al., 2024; Raffini et al., 2025). An advantage of studying *L. saxatilis* is that as a direct developer its lifetime dispersal is limited to a few meters (Janson 1983; Johannesson & Johannesson, 1995; Erlandsson et al., 1998; Westram et al., 2018), which should increase the propensity for local adaptation relative to over marine invertebrates (Sanford & Kelly, 2011). Two ecotypes have been extensively studied in Europe; the *Crab* and *Wave* ecotypes (Johannesson, 2003; Johannesson et al., 2024). The *Crab* ecotype inhabits sheltered boulder shores and is resistant to predation from crabs (Johannesson, 1986; Boulding et al., 2017), while the *Wave* ecotype has adapted to avoid dislodgement by strong wave action on exposed rocky shores that lack crabs (Larsson, 2021, pp. 59–80; Le Pennec et al., 2017). These ecotypes show strong phenotypic and genetic changes across sharp habitat transitions (Janson, 1982; Janson & Sundberg, 1983; Ravinet et al., 2016; Westram et al., 2018), which are paralleled by gradients in arrangement frequencies at 18 chromosomal inversions (Faria et al 2019; Morales et al., 2019; Westram et al., 2021; Koch et al., 2021, 2022; Hearn et al., 2022). While the *Crab*-*Wave* ecotype pair has received a lot of attention, other phenotypic varieties of *L. saxatilis* have been studied less, including the high spired *Brackish* ecotype (sometimes called ‘*tenebrosa’*; Reid, 1996) found in very sheltered habitats defined by soft sediments (Janson & Ward, 1985; Barnes, 1993; Johannesson and Johannesson, 1996). Earlier studies have also indicated vertical shore gradients in both genetic and phenotypic traits (Johannesson & Johannesson 1990; Morales et al 2019). While inversion arrangement frequencies commonly differ among ecotypes and shore levels, it has not been possible to identify the independent adaptive effects of each inversion although unusually strong differentiation and associations with adaptative phenotypes strongly support the contributions of some inversions (Morales et al., 2019; Koch et al., 2022). Analysing a broad set of phenotypic variables and inversion arrangement frequencies across all possible habitats can disentangle the role of individual inversions in local adaptation from the joint response of multiple inversions that we have observed over *Crab*-*Wave* hybrid zones (Westram et al., 2021). Hence, we used an unbiased sampling design to cover the full range of *L. saxatilis* habitats in an area in Sweden to contextualise the role chromosomal inversions have in the relationship between phenotypic, environmental, and genetic variation in a wider landscape.

## Materials and Methods

### Sampling snails

From April to November 2020, 200 intertidal sites were visited across an archipelago on the Swedish West Coast (Fig. 1A; Table S1) where the tidal range is small (∼30cm). Sampling sites were selected randomly, without prior information about habitats, from a 12x12km region which was subdivided into a grid of 100x100m cells. Cells containing only land or sea were excluded. 200 of the remaining cells were drawn from a sampling distribution, which was weighted (Pr(sample) = 0.9e^-5*dist/max.dist^ + 0.1) towards sites close to a well-described *Crab-Wave* hybrid zone (*Ängklåvbukten*; Faria et al., 2019; Westram et al., 2018, 2021). Snails were collected from the closest accessible shore to the centre of each cell. When no snail was found in a cell, it was recorded as a ‘No Snail’ site.

**Figure 1:**
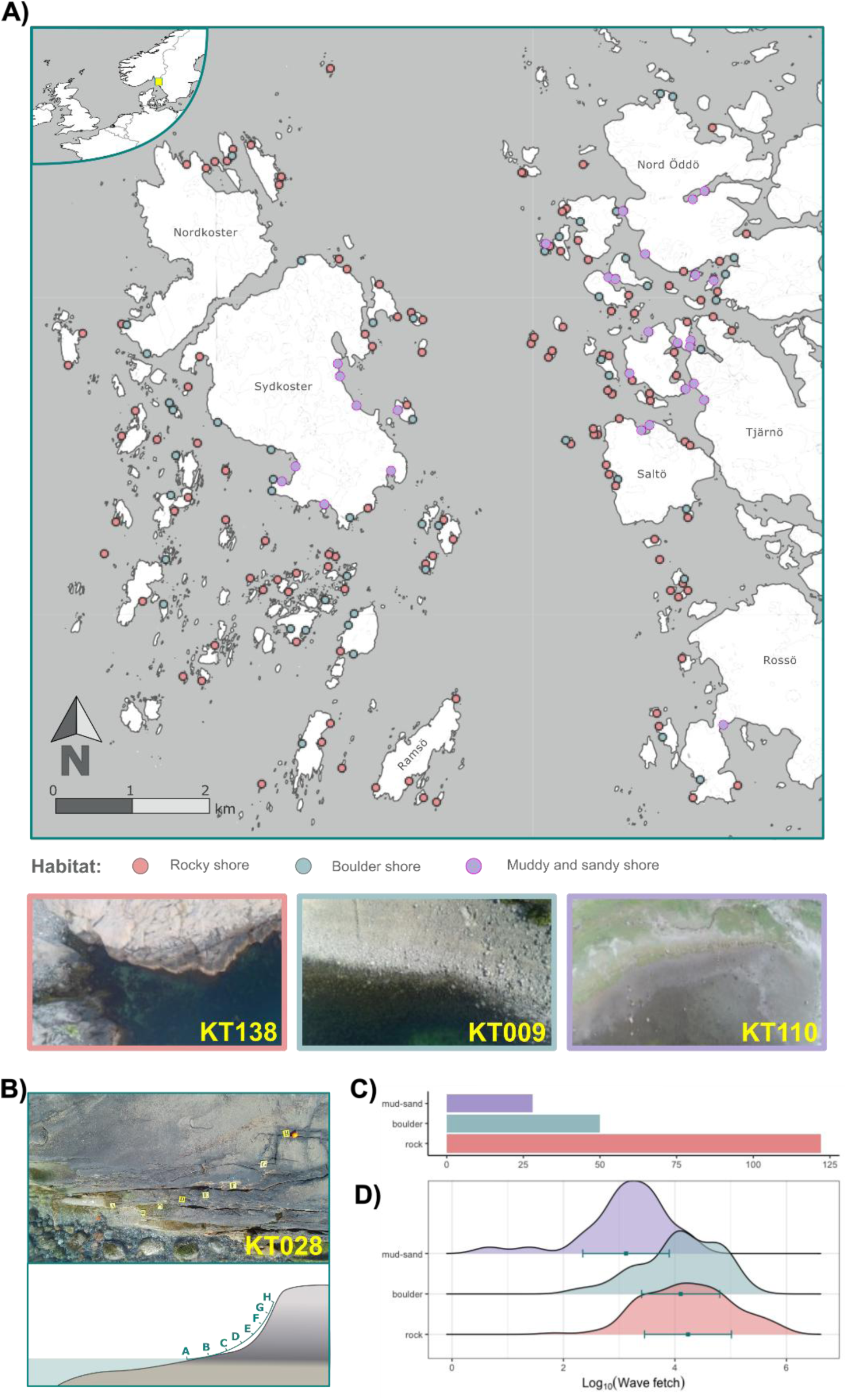
Sampling details. **A)** Map of all 200 sampling sites. Points are coloured by habitat type. Drone photos from three representative sites are below the map. The map is projected on the SWEREF99 CRS. Mapping data comes from the Swedish Land Survey (*Lantmäteriet*), accessed through the GeoData Extraction Tool (https://www.slu.se/en/subweb/library/use-the-library/search-and-find/find-digital-maps-and-geodata/). The insert map in the top left shows the sampling region’s location in the North Sea as a yellow square. **B)** Photo and simplified diagram of a sampling transect. **C)** Count of sites in each habitat type. **D)** Distributions for the log of wave fetch in each habitat type. Green dots and error bars represent the mean and standard deviation.

Snails were collected along a 7m transect that was set out along a shore height gradient at each site. A lead weighted rope was drawn from the lowest to the highest points of the snails’ distribution to define the transect (Fig. 1B). Often transects were not perpendicular to the shoreline, and in a few cases (e.g. skerries smaller than 7m, or smooth rock ledges) no transect was placed but snails were picked as far apart as possible. One snail was collected every meter along the transect, for a total of eight per site. The closest snail to each transect point was sampled. If no snail was found within 0.5m parallel or 1m perpendicular to the run of the transect from a sampling point, that point was considered empty. Snails shorter than 2.5mm were ignored, as they were unlikely to yield enough DNA for our genotyping approach. Each snail was labelled with the site ID (KT001 – KT200) and a letter corresponding to its position on the transect (i.e. ‘a’ was lowest and ‘h’ was highest on the shore). For sites without transects, snails were labelled in the order they were sampled.

The coordinates of each site were recorded at the bottom of the transect. Transect height was estimated using a yard stick with markings every 30cm. Heights were measured from sea level at the time of sampling to the highest point where snails were sampled. Measurements were subsequently adjusted to give heights relative to mean sea level, recorded by a nearby monitoring station (SMHI Sea Level data, Kungsvik).

### Measuring environments

A SwellPro SPRY+ drone was flown 10, 20 and 50m above each site to take photos of the transect, with its SONY 1/2.3’’ CMOS camera. Environmental features, which were used as proxies for major environmental gradients influencing local adaptation in *L. saxatilis* (e.g. wave exposure, predation, temperature stress, desiccation), were measured from these photos using ImageJ (Schneider et al., 2012). Before measurement, lens distortion was corrected with Adobe Lightroom’s default adjustment tool for SONY lenses. Sites were grouped into three broad habitat categories: rocky shores (**RS**), boulder shores (**BS**) and muddy or sandy shores (**MSS**), based on the prevailing habitat in each photo. Most environmental measurements were the same for each habitat type, with a few, such as rock fissures (RF), exclusive to a habitat type (Table 1; see supplementary methods1 for details). Additional observations of the environment were also taken while collecting snails. These included the presence of barnacles (*Semibalanus balanoides*) indicating stronger wave action, and the presence of brown algae *Fucus sp.* and *Ascophyllum nodosum* indicating less wave action and presence of predatory crabs hiding in the seaweed canopy (Lewis, 1972; Isæus, 2004).

**Table 1.**
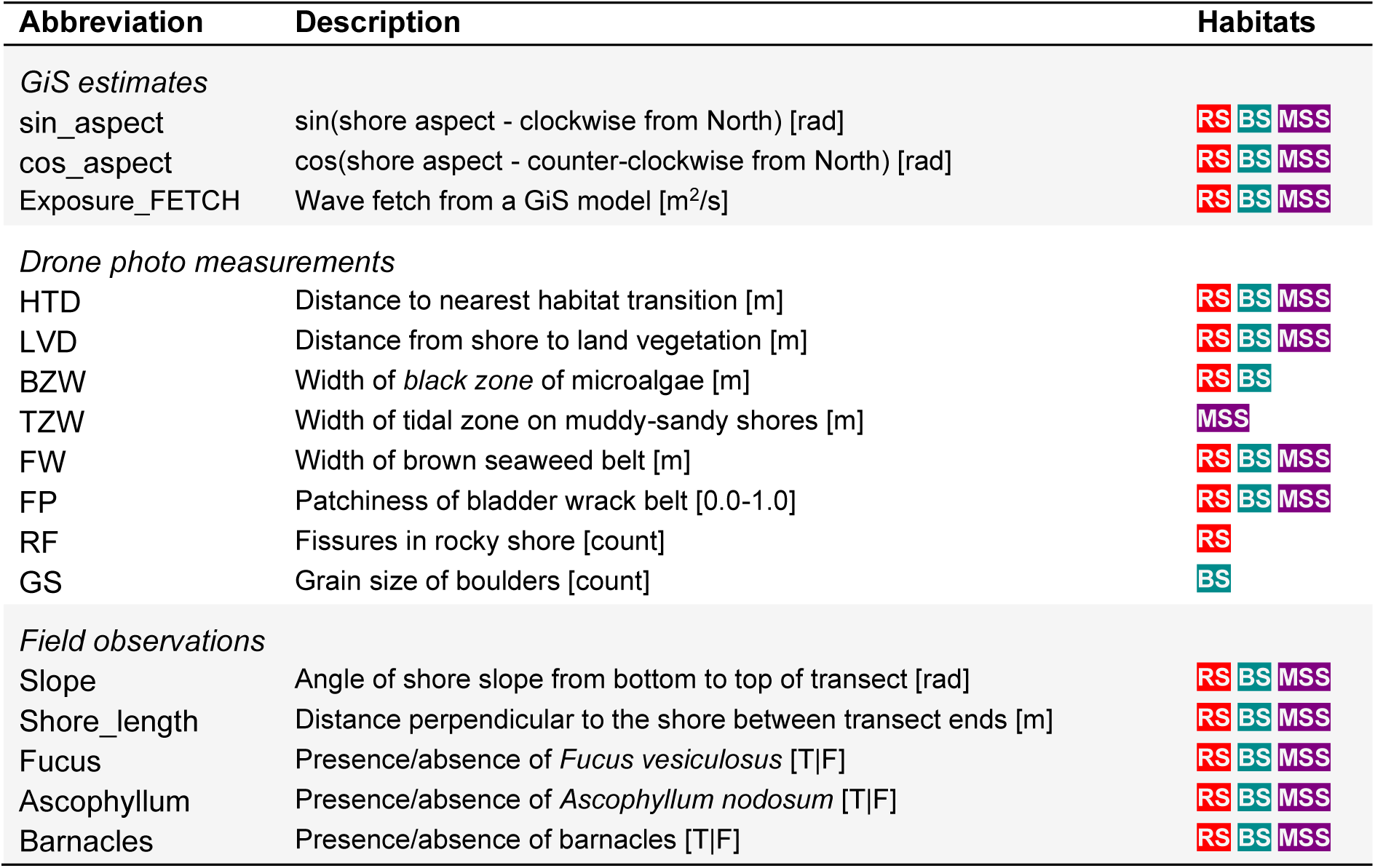
Abbreviations and descriptions of environmental variables. Units of measurement are indicated in square brackets at the end of each description. T|F represents a Boolean value. Habitat indicates which habitat classes each variable was measured in: **RS** = rocky shores, **BS** = boulder shores, and **MSS** = muddy and sandy shores. Drone photo measurements are explained in Supplementary methods 1.

Length measurements from drone photos were adjusted to correct for the slope of each shore. Slopes were estimated from the transect height and the horizontal distance from transect bottom to top of the shore, at each site. Length measurements perpendicular to the shore (i.e. LVD, BZW, TZW, and FW; Table 1), were divided by the cosine of the slope estimate. Two vertical cliff sites (KT195 & KT196) were not adjusted as they were photographed horizontally from the water.

Lastly, wave fetch and shore aspect were calculated in ArcGiS. Fetch was extracted by inputting site coordinates into a wave exposure model (Albertsson et al., 2006; Isæus, 2004). Shore aspect was calculated using the shore aspect tool in the *ArcMap* spatial analysis toolbox. Aspect scores were converted from degrees to radians in *R*. To avoid overestimating the differences between north facing shores (e.g. *R* calculates the differences between 350° and 10° as 340; it should be 20) aspect scores were split into the sin(aspect) moving clockwise from North and the cos(aspect) counterclockwise from North.

### Snail phenotyping and dissections

Snails were photographed in the lab, horizontally aligning the shell apex and columella. These photos were analysed for eight measurements of shell shape using the MatLab program **ShellShaper** (Larsson, 2020). Shell thickness was measured using a digital dial indicator (Neoteck 0.001–12.7mm range). The needle of the gauge was placed ∼1mm from the outer lip of the aperture, roughly a third of the way from the columella to the aperture suture.

Shells were broken after measurement, and the soft tissue of the snails was inspected. Sex and maturity were recorded. Maturity was determined by the presence of embryos in the brood pouch of females, or the presence of penial glands in mature males. Sex was determined by the presence or absence of a penis. Snails with neither a penis nor a brood pouch were recorded as juvenile females assuming that the brood pouch was too small to be detected. Each snail was then decapitated, just below the mantle tissue, and the head and foot were stored in 95% ethanol for DNA extraction.

### Dimension reduction of phenotypic and environmental variables

Environmental variables were reduced to a few orthogonal axes for analysis using the R package **PCAmixdata** (Chavent et al., 2022). This package combines PCA of continuous variables (e.g. BZW, LVD, HTD, Shore_length; Table 1) with multiple correspondence analysis (MCA) of categorical variables (e.g. barnacles, *Ascophyllum*, *Fucus*; Table 1). As the reorientated axes of environmental variation are not strictly principal components, they will be referred to as dimensions in this text (e.g., Dim1 – 3). PCAmix analysis was performed separately for each of the three main habitat types. Eigenvalues were inspected to determine the number of dimensions to return by comparing with a broken stick model. The influence each original variable had on a dimension (factor loading) was determined by inspecting the eigenvectors (Fig. 2B).

**Figure 2:**
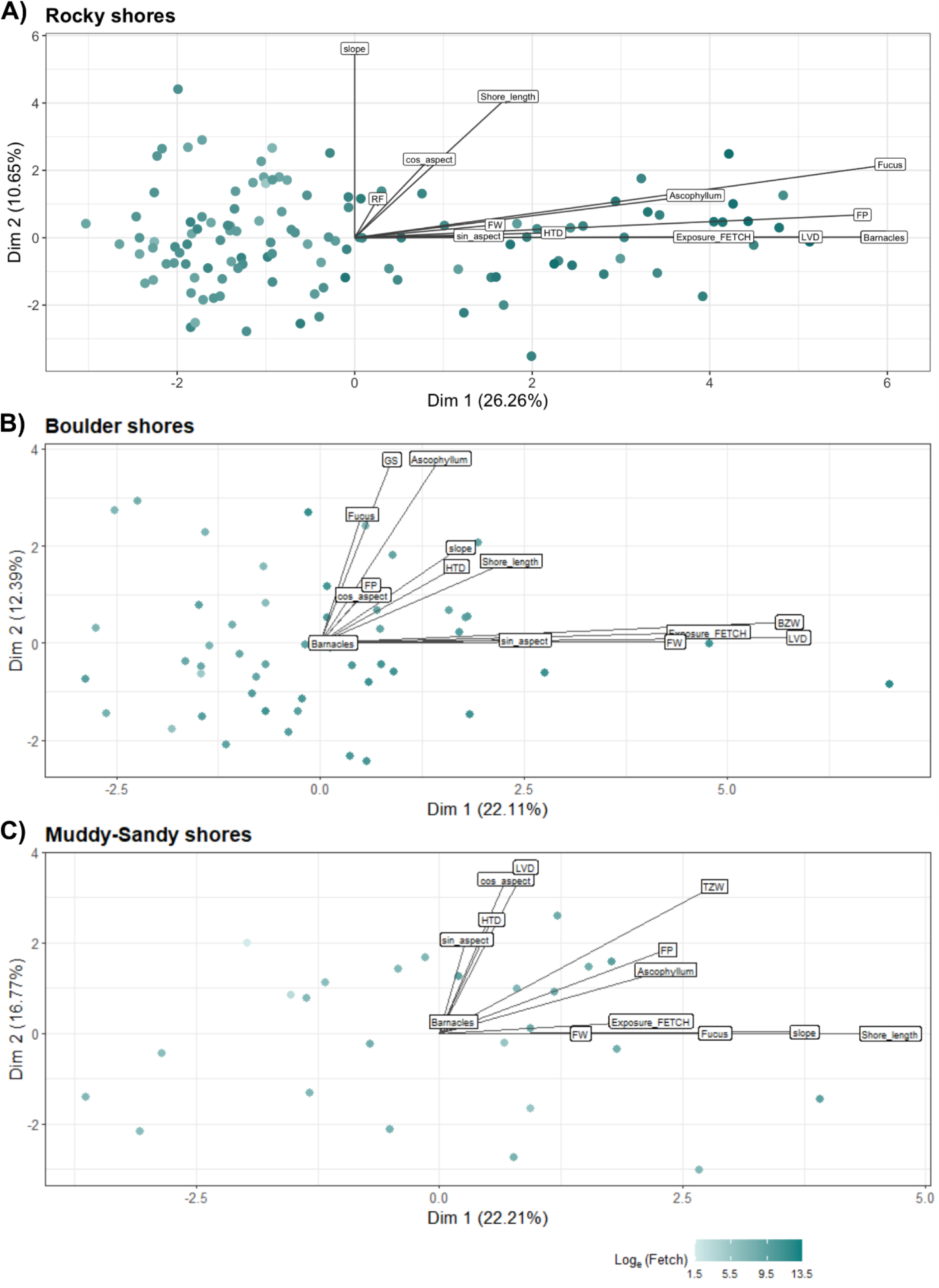
PCAmix results for **A)l** rocky shores, **B)** boulder shores, and **C)** muddy-sandy shores. Points indicate sites, coloured by log-fetch as a measure of wave exposure. Labels are the factor loadings of each environmental variable. Descriptions of the variables are in Table 1. Further dimensions, distribution plots and scree plots for each habitat type are in the supplementary materials (Fig. S1-3).

Phenotypic variables, excluding shell length (Table 2), were reduced to a few orthogonal axes, with PCA implemented with the R function *prcomp()*. Phenotypes were standardised before PCA; first all distance measurements (avg_thick_mm, r0, z0 and a0) were divided by the total shell length to correct shape for snail size. Eccentricity was then divided by the aperture radius (a0). These adjustments were followed by scaling and centring all variables with the R function *scale()* before running *prcomp()*. The resulting eigenvalues were evaluated in the same way as the environmental dimensions. The full R scripts for dimension reductions are available on GitHub (https://github.com/ja-Reeve/Koster_data_analysis/dimension_reduction).

**Table 2.** Abbreviations and descriptions of each phenotypic variable. Units of measurement are indicated at the end of each description. Variables used in PCA are separated from co-factors which were included in Bayesian regression analysis. ShellShaper variables are explained in greater detail in Larsson et al. (2020); note that in Larsson et al. (2020) z0 was renamed as h0.

| Abbreviation | Description | Source |
| --- | --- | --- |
| *shellLength* | *Distance from bottom of shell to spire apex [mm]* | *ShellShaper* |
| *Included in phenotype PCA* |  |  |
| z0 | Distance from centre of aperture to spire apex [mm] | ShellShaper |
| r0 | Distance from centre of aperture to spire meridian [mm] | ShellShaper |
| a0 | Radius of aperture [mm] | ShellShaper |
| eccentricity | Eccentricity of aperture [0 = circle; > 0 more ovular] | ShellShaper |
| apAngle | Angle at the aperture join [°] | ShellShaper |
| gw | Growth parameter for shell width: $gw = \ln\left(\frac{r0+a0}{w1}\right)/(2\pi)$ | ShellShaper |
| gh | Growth parameter for shell height: $gh = \ln\left(\frac{z0}{w1}\right)/(2\pi)$ | ShellShaper |
| avg_thick_mm | Average thickness of shell at lip of aperture [mm] | Measurement |
| *Co-factors in analysis* |  |  |
| sex | Male or female | Dissection |
| maturity | Adult or juvenile | Dissection |

### Inferring ecotypes

To facilitate comparisons between the phenotypic and environmental axes derived here and previous work, snails from 83 sites were assigned to the *Crab* (48) or *Wave* (35) ecotype based on habitat descriptions from previous studies (Johannesson et al., 2010; Westram et al., 2021), and subjective inferences of habitat type from drone photos and morphology in shell photos. These included two sites (KT007 and KT009) from a *Crab-Wave* hybrid zone (*Ängklåvbukten*) where phenotypic and genomic variation have earlier been analysed in detail (Janson & Sundberg 1983; Westram et al., 2018; Faria et al., 2019).

### SNP genotyping and inversion genotype inferences

Snails were genotyped to determine the contributions of inversions and the genetic background to phenotypic variation. Genotyping was a two-step process, with 98 snails whole genome sequenced (WGS) to design a SNP panel with which to genotype the remaining 1,389 snails. First DNA was extracted from snail foot tissue using a CTAB-chloroform protocol (Panova et al. 2014). Libraries were prepared and sequenced using DNBSEQ to approximately 5X coverage at BGI genomics, Hong Kong. Reads were then processed into SNPs using a BWA-bcftools pipeline. See supplementary method 2 for details on DNA extractions and WGS processing.

A SNP panel was developed from the WGS data to genotype the remaining snails. The panel included 102 SNPs that were strongly associated with inversions, and a subset of 198 SNPs was sampled from the colinear genome, which was called the “genetic background”. Supplementary method 3 provides more details on SNP panel design.

Genotypes for each inversion were assigned based on a PCA of all SNP markers within the inverted region, following the approach detailed in Faria et al. (2019) and Reeve et al. (2024). PCAs were run following the protocol for designing the SNP panel (see supplementary method 3). In a few cases SNPs that had low PCA loadings had to be removed to differentiate PCA clusters.

### Isolation by distance

Isolation by distance was assessed among all snails by comparing Euclidian spatial distances to genetic distance. Spatial distances were calculated among sites with the **QGIS** (v3.16, 2020) distance matrix tool using the default parameters. Patterns of isolation by distance are expected across the sampling region, as the lack of a pelagic larval stage makes *L. saxatilis* dispersal distances shorter than many other marine invertebrates (Johannesson & Johannesson, 1995; Sanford & Kelly, 2011). Note, only straight-line distances were used, which may not perfectly represent snail dispersal. Genetic distance among samples was calculated using Edward’s distance, implemented through the **R** function *dist.genpop()* in the *adegenet* package (Jombart, 2008; Jombart & Ahmed, 2011). Only genetic background SNPs were used to calculate genetic distance. The correlation between spatial distances and genetic distances was assessed using a Mantel Test (Mantel, 1967). This relationship was plotted as a density matrix using a 2D kernel density estimation from the **R** package *MASS* (Venables & Ripley, 2002). A linear model was superimposed to show the average trend across sites. In addition, spatial PCA was used to determine if there was geographic clustering across the sampling region (Jombart et al., 2008). Only one, large-scale spatial axis was retained (sPC1).

### Modelling relationships between environment, phenotype and inversions

Bayesian regression models were run using the R package *brms* (Bürkner, 2017) to investigate the relationships between environmental variables and inversion frequencies, and their associations with phenotypes. Each analysis asked a different question:

**Pheno∼Inv:** to what extent do the arrangement frequencies of each inversion explain each axis of phenotypic variation?

**Pheno∼Env:** how does phenotypic variation relate to environmental variation?

**Inv∼Env:** does environmental variation help to explain inversion arrangement frequencies? The full numbers and sets of parameters included in each analysis are explained in Table 3. Sex and maturity were included as individual-level effects in every model. Genetic covariance among samples and sampling date among sites were treated as group-level random effects (Bürkner, 2017), to control for geographic and seasonal structure, respectively. Sex, maturity, genetic covariance, and sampling date are collectively referred to as co-factors. The height on shore for each snail was considered as an additional co-factor but ultimately dropped as it had limited effects in **Pheno∼Env** models (supplementary method 4). Both the **Pheno∼Env** and **Inv∼Env** models were run separately within each of the three main habitat types and for an among-habitat-class model. When the complex inversion LGC6.1/2 (probably two overlapping inversions resulting in three segregating arrangements; Faria et al., 2019) was a response variable, it was split into two **Inv∼Env** models; ‘LGC6.1/2’ compared the arrangement more common in *Crab* to the two alternative arrangements combined, and ‘LGC6.1/2b’ compared the frequencies of the two alternative arrangements. Inversions that have been previously associated with ecotype formation (LGC6.1/2, LGC14.1/2 and LGC17.1) and shore height (LGC9.1 and LGC12.2; supplementary method 4) were predicted to have the strongest associations in the **Pheno∼Inv** and **Inv∼Env** models.

**Table 3.**
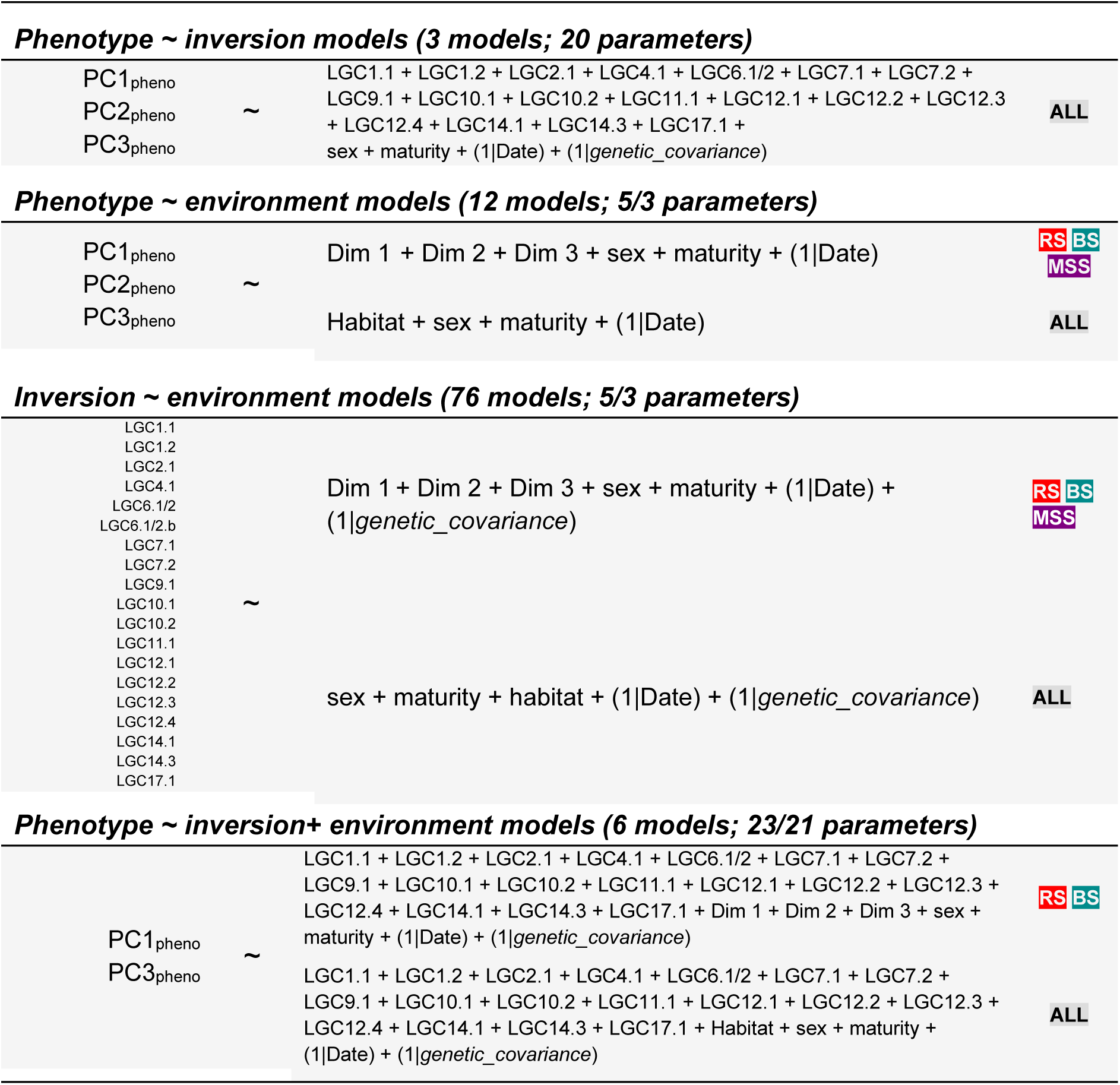
Overview of Bayesian regression models tested in brms. Response variables are shown in the left column. Titles indicate the analysis performed. The middle column shows the parameters tested in each model. Dim X = environmental PCAmix results. LGCx.y are the inversions. Right column indicates the different subsets of the data each model was tested on: **RS** = rocky shores, **BS** = boulder shores, **MSS** = muddy and sandy shores, and **ALL** = full dataset. PCX_pheno_ = phenotypic PCA results. Date = sampling date, which was treated as a random factor. *genetic_covariance* = a genetic covariance matrix among all samples, which was treated as a random factor.

To test the independent effects of environmental dimensions and inversion arrangements on phenotypic variation (i.e., phenotypic plasticity, effects of adaptive loci outside inversions and confounded variation between inversions and environment), models from the first round of Bayesian regressions were used to build **Pheno∼Inv+Env** models. Muddy-sandy shores were not considered in the **Pheno∼Inv+Env** models because the sample size was small and model fits were poor (based on ΔLOOIC) in the previous models. Six models were run in this final round, considering the effects of inversions and environmental variation on PC1_pheno_ and PC2_pheno_ within, and between, rocky shores and boulder shores.

A two-tiered approach was used to select informative terms from all models. Firstly, the overall fit of the model was evaluated with ΔLOOIC scores (Vehtari et al., 2015). These were evaluated like AIC, comparing each tested model to a null model with co-factors as predictors (i.e., LOOIC_Tested.model_ – LOOIC_Null.model_). Models with ΔLOOIC < -2 x standard error were retained. A secondary filter (ΔLOOIC < -4 retained) was set to remove models with little explanatory power that still might have passed the standard error filter. Informative terms were selected from retained models when their 95% posterior credibility intervals did not overlap 0. Separate models were built for each habitat class, plus an among-habitat model.

### Variance decomposition

Proportions of phenotypic variance explained by each variable were estimated using Bayes R^2^ (Gelman et al., 2018) for a series of sub-models. Variables were grouped into four major components: the variance explained by (i) inversion arrangements, (ii) environmental effects, (iii) the genetic background, and (iv) the co-factors sex, maturity and sampling date. Equations and a more detailed description of variance decomposition can be found in supplementary method 5.

## Results

### Distribution of snails and environmental variation

In total, 1,488 snails were collected from 196 of 200 sampled sites, the remaining four sites had no snails. Sites with snails represented an unexpectedly wide array of habitats (Fig. 1A); from snails living in the grass under trees beside a sheltered muddy lagoon (KT145) to heavily wave-exposed rocks facing the open sea (KT138). Only sandy shores without gravel and a smooth rocky cliff without crevices were free of snails. Fewer than eight snails were collected from 53 sites (Table S1), sometimes due to species misclassifications (mostly juvenile *Littorina littorea*), but more often related to low snail densities in sheltered rocky sites. Wave exposure distributions (measured as fetch) overlapped among the three habitat classes (rocky, boulder and muddy-sandy shores; Fig. 1D). Rocky shores were more wave exposed on average, but much of this distribution overlapped with boulder shores (Fig. 1D).

Rocky shores were the most common sampling habitat (Fig. 1C), representing the full spectrum of wave fetch measured for this region (Fig. 1D), along with a wide array of other components of environmental variation summarised with PCAmix. The first dimension (Dim1_RS_) for rocky shores was loaded with terms related to wave exposure such as fetch, distance from the sea to land vegetation and presence of barnacles (Fig. 2A-B). Dim1_RS_ explained 26% of all environmental variation (Fig. 2E). Its distribution was broad (skewness = 0.85; kurtosis = 2.53), running from very sheltered to very exposed sites (Fig. 2C), but it was not significantly different from a unimodal distribution (Hartigan’s dip test; D = 0.02; P = 0.99). Dim2_RS_ was loaded with variables more related to shore topography, like slope, shore length and aspect (Fig. 2B). Factor loadings for other dimensions could not be interpreted in a simple way.

Biological interpretations of environmental loadings were less clear for boulder shores and muddy-sandy shores. On boulder shores, Dim1_BS_ was loaded with terms related to wave exposure (Fig. S2). Factor loadings provided no clear interpretation for muddy-sandy shores (Fig. S3). The scree plot was flatter, with earlier dimensions containing less variance than expected (Fig. S3E). In fact, a common set of environmental measurements across habitats showed that muddy-sandy shores were more homogeneous than rocky shores (Fig. S4).

### Phenotypic continuity

Phenotypes did not fall into discrete groups. No clusters of snails could be seen on any principal component (Fig. 3B; Fig. S5E) or along the distribution of shell lengths (Fig. 4A). PC1_Pheno_ explained 43% of the variation in shape phenotypes, which was loaded for factors related to whorl growth, spire height and aperture size (Fig. S5C). However, PC1_Pheno_ was not normally distributed (Fig. S5D), with variation on this axis being flatter than a normal distribution (kurtosis = 2.74) and with a slight negative skew (skewness = -0.04). This could reflect both ecotype differentiation and sexual dimorphism. There was a significant difference in average PC1_Pheno_ between sexes (male *x̄* = −0.15, *σ* = 1.69; female *x̄* = 0.11, *σ* = 1.94; t = -2.77, df = 1414, P = 0.006). PC2_Pheno_ was loaded for factors related to the shell aperture. This axis should be interpreted cautiously as aperture angle is poorly defined when the aperture is close to being circular rather than elliptical (Jenny Larsson, personal communication). Principal components beyond PC2_Pheno_ were not considered as they contained less variance than expected by chance under a broken-stick distribution (Fig. S5F).

**Figure 3:**
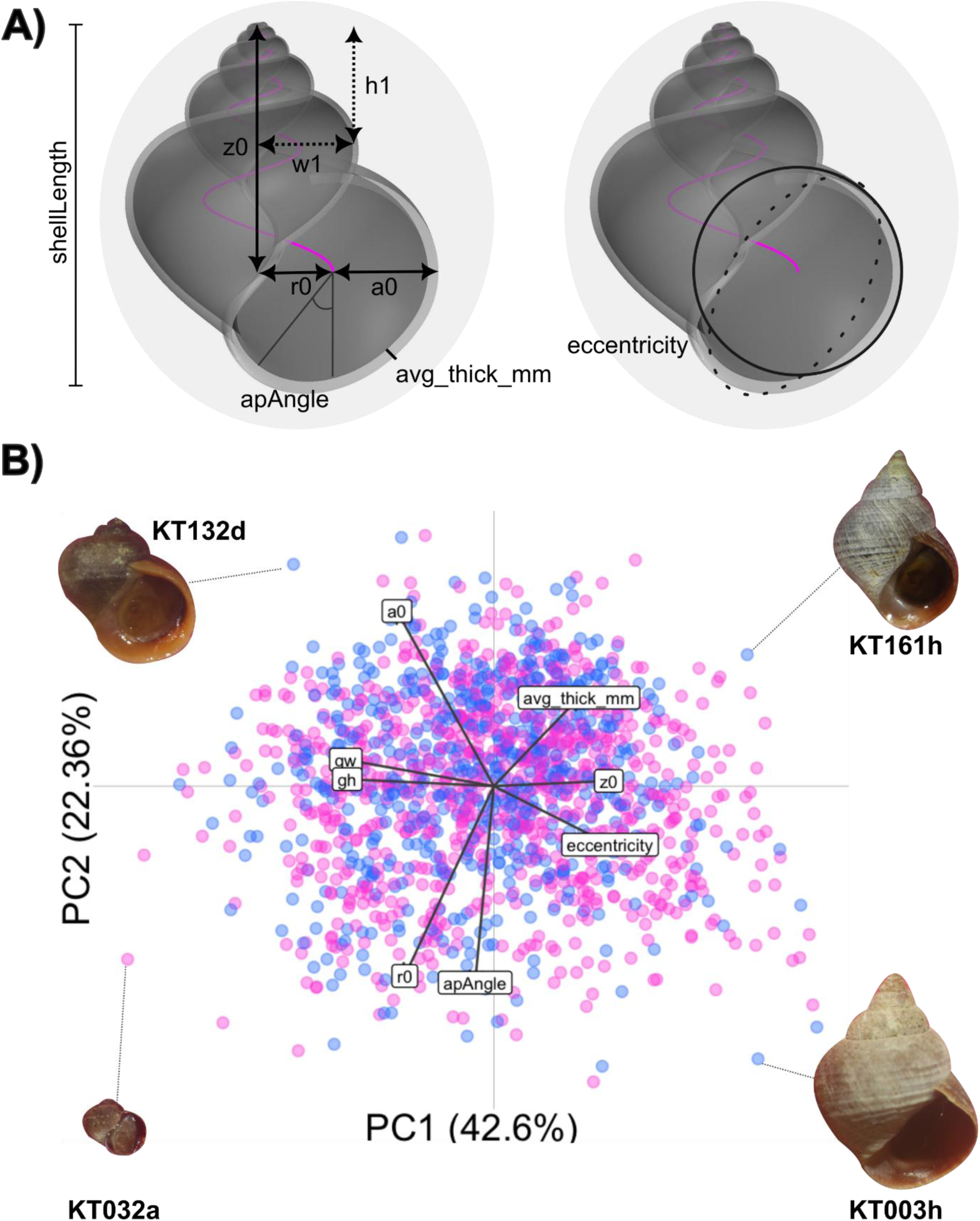
PCA results for all 1,488 snails. **A)** Diagram of a ShellShaper model indicating measurements of shells. The ‘growth parameters’ gw and gh are calculated from the additional measurements w1 and h1: *gw* = ln 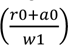 /(2*π*) and *g*ℎ = ln 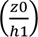 /(2*π*). **B)** PC1 vs. PC2 scatter plot. Points are coloured by sex (pink = female; blue = male). Labels are the factor loadings of each phenotypic variable. Four shell photos surrounding the plot. These were selected as the extreme edges of the principal components. Factor loadings for, PC distributions and scree plots are in Fig. S4.

**Figure 4:**
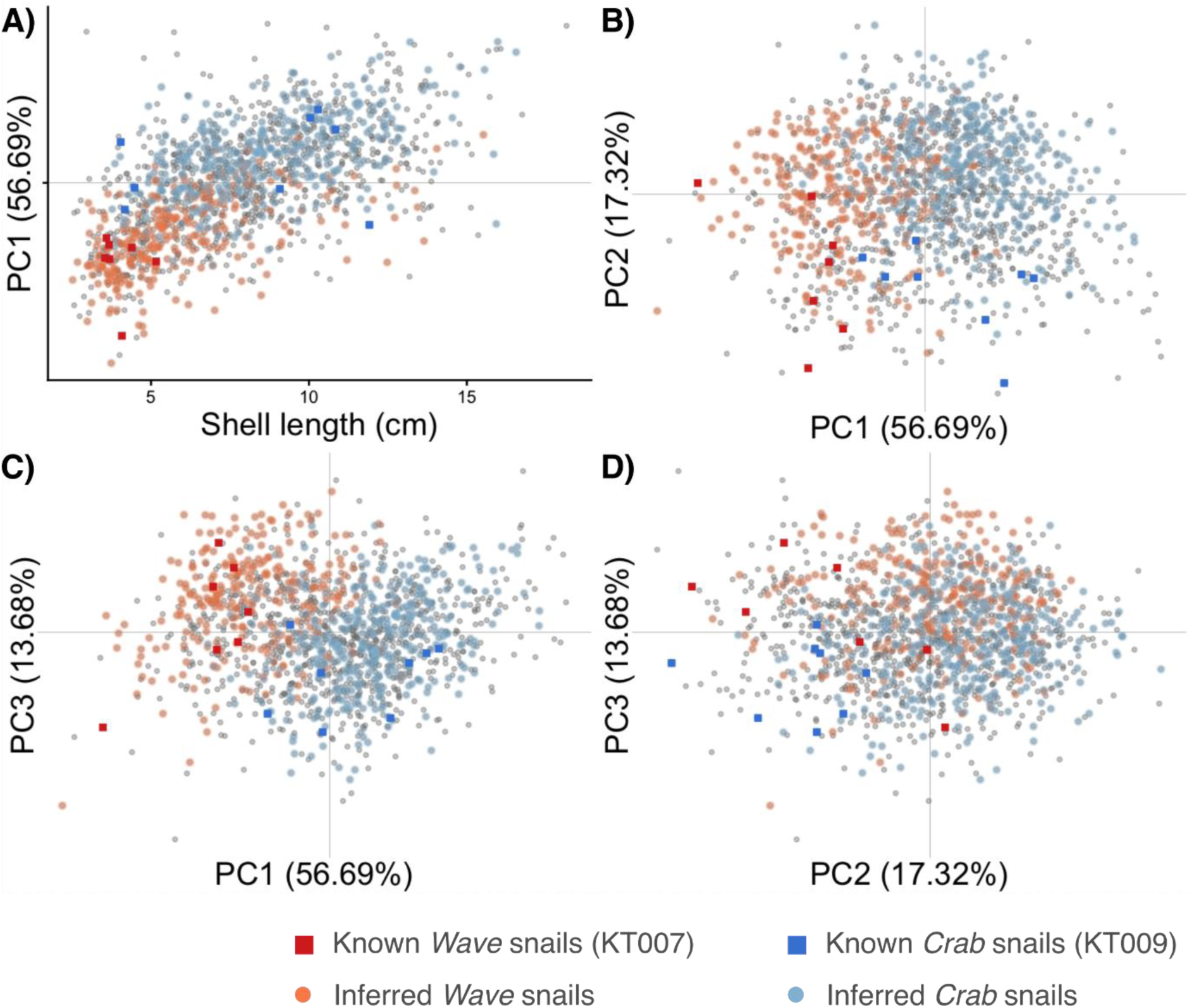
Scatterplots of phenotypic variables, colouring the snails which correspond to *Crab* (blue) and *Wave* (red) ecotypes. **A)** shell length vs. PC1_Pheno_. **B)** PC1_Pheno_ vs. PC2_Pheno_, **C)** PC1_Pheno_ vs. PC3_Pheno_. **D)** PC2_Pheno_ vs. PC3_Pheno_. Squares indicate snails from sites that were previously studied for ecotype divergence (Janson & Sundberg, 1983; Koch et al., 2021, 2022; Westram et al., 2018, 2021). Lighter shaded dots indicate snails with inferred ecotypes. Grey points representing snails from other sites.

The phenotypic principal components distributed the snails assigned to the *Crab* and *Wave* ecotypes towards opposite ends of a continuum. Snails from known ecotype sites separated out along PC1_Pheno_, with a minor overlap. A similar pattern was seen along PC1_Pheno_ for snails with inferred ecotype labels (Fig. 4), with a significant difference in means despite considerable overlap (*Crab x̄* = 0.92, *σ* = 1.33; *Wave x̄* = −1.73, *σ* = 1.29; Welch’s t = 25.11, df = 578.08, P < 2.2x10^-6^). *Crab* snails were generally longer than *Wave* snails (*Crab x̄* = 8.71*mm*, *σ* = 2.57*mm*; *Wave x̄* = 5.98*mm*, *σ* = 2.38*mm*; Welch’s t = 13.76, df = 592.25, P < 2.2x10^-6^; Fig. 4A). The phenotypic gradient was larger than the ecotypes, including snails from both extreme and intermediate phenotypes from sites which were not assigned to an ecotype (Fig. 4).

### Isolation by distance

Background SNP genotypes of snails from different sampling sites revealed fine-scale isolation by distance. Genetic and spatial distances were only weakly correlated (Mantel test: ⍴ = 0.14; replicates = 999; P = 0.001). The slope of the relationship between log-spatial distance and genetic distance was very shallow (β = 0.003). When projected onto the untransformed spatial distance axis (Fig. S6) there was little increase in genetic distance for sites >2km apart, which is expected as mean dispersal in *L. saxatilis* is only 1-3m per generation and isolation effects are already saturated at short distances due to the lack of a pelagic larval stage (Janson 1983; Johannesson & Johannesson, 1995; Erlandsson et al., 1998; Westram et al., 2018). Spatial PCA indicated that background genetic structure may be related to wave exposure, with more windward and less enclosed sites tending to group together on the one retained spatial PC (Fig. S7). This suggests either that some putatively neutral loci are influenced by selection, maybe due to linkage with inversions, or that gene flow is stronger between snails from similar habitats (i.e., isolation by environment; Wang & Bradburd, 2014).

### Regression analysis 1.1: effects of sex and maturity

Variation in phenotypes, environments, and inversions were compared in a series of Bayesian regression models. Sets of factors were compared in pairs (**Pheno∼Env, Pheno∼Inv**, and **Inv∼Env**), followed by a combined analysis of all factors which controlled for confounding of effects on phenotypes between inversion arrangement frequencies and environmental variables (**Pheno∼Inv+Env**).

Sex, maturity and sampling date were considered as co-factors in these models. Sex differences were associated with shell length and weakly associated with the phenotypic PCs (Fig. S8A-C). For example, in the among-habitat models (which included all samples), males were on average 1.03 mm (95% credibility intervals: 0.80 – 1.26) shorter than females; while the average difference in PC1_Pheno_ between females and males was -0.52 (−0.67 - -0.38), representing proportionally shorter and broader spires in females. The average difference in PC2_Pheno_ between sexes of 0.16 (0.05 – 0.26) suggests females have apertures more centrally located relative to the spire axis. Four inversions were correlated with sex (Fig. S8D), including two inversions on LG12 that are linked with a sex determining locus for the *Crab* ecotype (Ghane, 2022; Hearn et al., 2022).

Snail maturity influenced shell length in the **Pheno∼Env** model as expected (Fig. S8A), but also arrangement frequencies of eight inversions in the **Inv∼Env** models (Fig. S8D). On average, adults were 2.67 mm (2.40 – 2.93) longer than juveniles and shell shape differed with snail maturity (Fig. S8), supporting existing evidence of allometric growth (Hollander et al., 2006a). Maturity was most strongly associated with arrangement frequencies in two inversions (LGC14.1/2, and LGC17.1) in the full dataset, and smaller changes in arrangement frequencies of six other inversions (LGC4.1, LGC10.1, LGC11.1, LGC12.1, LGC12.2, and LGC12.4; Fig. S8D). These patterns may represent snails that mature at smaller sizes in exposed habitats, which has confounded environmental and maturity effects with inversion arrangement frequencies.

### Regression analysis 1.2: phenotype and environment

The strongest phenotypic differences among habitat classes were for shell length and PC1_Pheno_ (Fig. 5A, Fig. S8). Compared with boulder shores, snails were on average 1.99 mm (1.68 – 2.31) shorter on rocky shores and 1.62 mm (1.13 – 2.10) shorter on muddy-sandy shores. There were also smaller, yet informative differences for average PC1_Pheno_ between boulder and rocky shore snails (PC1_Pheno,BS_ – PC1_Pheno,RS_ = -1.59 [-1.39 – -1.79]) and PC2_Pheno_ (PC2_Pheno,BS_ – PC2_Pheno,RS_ = -0.30 [-0.18 – -0.43]). Boulder and muddy-sandy shores only had informative differences along PC1_Pheno_ (PC1_Pheno,BS_ – PC1_Pheno,MSS_ = -0.77 [-0.47 – - 1.07]). Generally, **Pheno∼Env** models only weakly improved upon the null models, explaining 8% of phenotypic variance at most, and <2% for most PC2_Pheno_ models (Fig. 5B).

**Figure 5:**
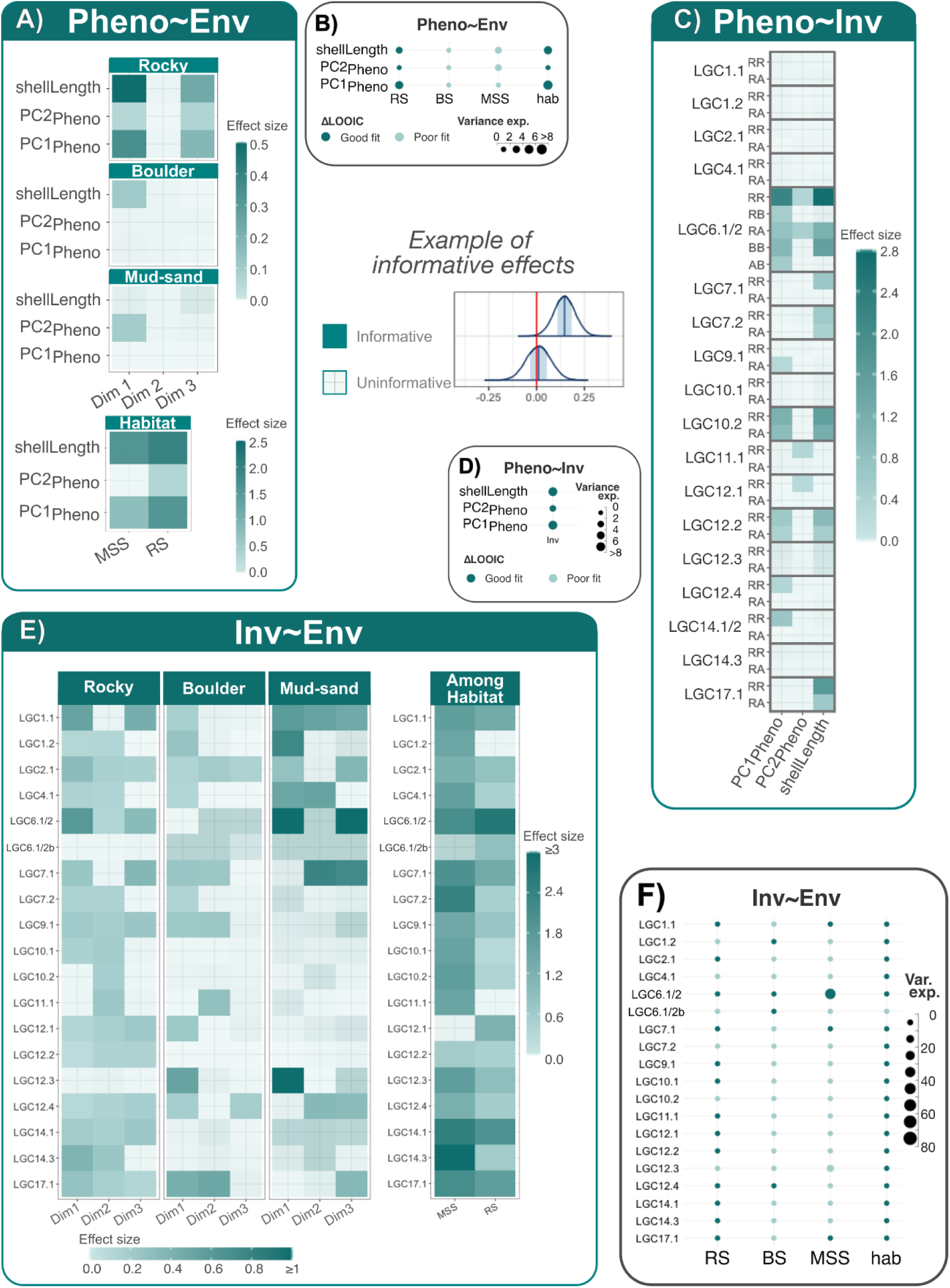
Bayesian regression model results. **A)** Tile plot summary of **Pheno∼Env** models including the effects of the environmental PCAmix axes (Dim1-3), the differences between boulder and muddy sandy shores (**MSS**), and between boulder and rocky shores (**RS**). Filled tiles are informative (based on posterior distributions; example shown to the right). Darker shades indicate stronger effect sizes. Tiles with a cross were uninformative. **B)** Dot plot summary showing the percentage of variance explained (Bayes R^2^) and ΔLOOIC scores for these **Pheno∼Env** models. **C)** Tile plot of **Pheno∼Inv** models. **D)** Dot plot summary of **Pheno∼Inv** models. **E)** Tile plot summary of **Inv∼Env** models including PCAmix axes (Dim1-3) and the differences among habitats. LGC6.1/2b was the model comparing the two alternative arrangements of this inversion. **F)** Dot plot summary of **Inv∼Env** models. Effects of co-factors (Fig. S8) and posterior distributions (Fig. S9-S11) are presented in the supplementary materials.

The first dimension of environmental variation (Dim1) in each habitat class had the strongest associations with phenotype (Fig. 5A, Fig. S9). These associations were strongest between Dim1 and shell length, especially within rocky shores. This suggests that shell sizes decrease with exposure within rocky shores, and to a lesser extent increase within boulder shores, while sizes in muddy-sandy shores were less variable. Dim3_RS_ was also predictive for all three phenotypic variables in rocky shores (Fig. 5A, Fig. S9), but a biological interpretation for these associations is not clear.

Within muddy-sandy shores Dim1_MSS_ influenced PC2_Pheno_, however the variance explained was low for most models (Fig. 5B). Wide credibility intervals reflect the low power of this data set, as few muddy-sandy shores were visited in our randomised sampling (Fig. 1C). We observed these shores to be more homogeneous environments than shores of the other habitat types, thus we expect a reduced pressure for within-habitat local adaptation in the muddy-sandy habitat.

Snail height on shore was tested as an additional environmental variable but had minimal impact on **Pheno∼Env** models (see supplementary method 4), explaining at most 2% of phenotypic variance (Table S2). Furthermore, shore height models did not improve model performances (evaluated with LOOIC differences) over the **Pheno∼Env** models (Table S2). To reduce total parameter counts, and prevent overfitting, snail height on shore was excluded from the main analysis.

### Regression analysis 1.3: phenotypes and inversions

Phenotypic variation was associated with over half (11 of 18) of the inversions in at least one **Pheno∼Inv** model (Fig. 5C). Six inversions influenced shell length and PC1_Pheno_, three of which (LGC6.1/2, LGC10.2 and LGC12.2) were jointly associated with both phenotypic axes. Only three inversions affected PC2_Pheno_, two of which were exclusive (LGC11.1 and LGC12.1). Only LGC6.1/2 had a shared influence on every phenotypic axis (Fig. 5C, Fig. S10). Candidate inversions for ecotype divergence generally had effects in the **Pheno∼Inv** models, with strong associations with LGC6.1/2 and LCG17.1, and moderate effects of LGC14.1/2 on PC1_Pheno_. Conversely, LGC10.2 had strong associations, but was only a weak candidate for ecotype divergence in previous studies (Morales et al., 2019; Westram et al., 2021; Koch et al., 2022; Reeve et al., 2024; Raffini et al., 2025).

Two potential effects came from LGC12.3 and LGC14.3. These two inversions yielded large but uninformative estimates, i.e. centred away from zero but with very wide posterior distributions of slope (Fig. S10). This higher uncertainty stems from each inversion having a low minor arrangement frequency (*overall alternative arrangement frequencies*: p_LGC12.3_ = 0.03; p_LGC14.3_ = 0.05; *average across inversions* = 0.25 ± 0.06), resulting in low power.

### Regression analysis 1.4: inversions and environment

Arrangement frequencies varied among habitat classes for every inversion, with the largest differences between boulder and muddy-sandy shores (Fig. 5E, Fig. S11). Within habitats, environmental associations were strongest among rocky shores, with all inversions except LGC12.3 and LGC6.1/2b (the comparison of alternative arrangements in this complex inversion) having informative relationships with at least one axis of environmental variance. LGC12.3 was possibly associated with Dim2_RS_, however the alternative arrangement was rare in rocky shores (p_LGC12.3,RS_ = 0.03) leading to a wide posterior distribution (Fig. S11), and thus little explanatory power for this model (Fig. 5F). In contrast, both alternative arrangements of LGC6.1/2b were common (p_LGC6.1/2,**A**,RS_ = 0.18; p_LGC6.1/2,**B**,RS_ = 0.30), and posterior distributions of effect sizes were relatively narrow (Fig. S11). Given the strong effects on LGC6.1/2, the lack of informative effects on LGC6.1/2b, indicates that both alternative arrangements of this complex inversion (with three arrangements) have similar habitat-dependent fitness that together is different from the **R** arrangement. Further tests were run on LGC9.1 and LGC12.2 to investigate the potential effects of adding snail height on shore to selected **Inv∼Env** models, but this added term contributed at most 6% to the variance explained by the model (supplementary method 4).

Boulder shores had fewer associations between inversions and environment, but arrangement frequencies for two thirds of the inversions (12 of 18) were influenced by environmental variation (Fig. 5E). There were weak associations between environmental variables and LGC6.1/2, but these were uninformative with wide posterior distributions (Fig. S11). This uncertainty stems from the alternative arrangements of LGC6.1/2 being rare in boulder shores (p_LGC6.1/2,**R**,BS_ = 0.95; p_LGC6.1/2,**A**,BS_ = 0.01; p_LGC6.1/2,**B**,BS_ = 0.04).

Muddy-sandy shores had many clear links between environment and inversion arrangement frequencies (9 of 18), however as mentioned above for **Pheno∼Env** models (Fig. 5E, Fig. S10), muddy-sandy shore models generally had greater uncertainty due to lower sample sizes. Although muddy-sandy shores also had the most variance explained of any **Inv∼Env** model (Fig. 5F), and the largest effect sizes for LGC6.1/2 and LGC12.3 (Fig. 5E, Fig. S11), these results should be treated cautiously, as the posterior distributions were wide (Fig. S11) and Pareto-K scores indicated that the **Inv∼Env** models did not explain all data points well.

### Regression analysis 2.1: joint effects of environment and inversions on phenotype

**Pheno∼Inv+Env** models were run to disentangle the effects of environment on phenotypes from the variance explained by individual inversions. Muddy-sandy shore models were analysed, but the results were omitted from the main text, as the Bayesian models did not reach convergence after 40,000 iterations (see Fig. S12 for the posterior distributions). This lack of convergence likely was caused by small the number of samples from muddy-sandy shores (n = 187).

Phenotype∼environment effects were mostly consistent between **Pheno∼Inv+Env** (Fig. 6A) and **Pheno∼Env** models (Fig. 5A). The two strongest associations with Dim1_RS_ were still present in the **Pheno∼Inv+Env** models (shellLength and PC1_Pheno_), but with smaller effect sizes. Among habitat effects were also similar in the new models (Fig. 6A). In contrast, the associations with Dim3_RS_ were non-informative in the **Pheno∼Inv+Env** models (Fig. S8; Fig. S12). The strong environmental effects shown in the **Pheno∼Env** models were likely confounded with inversion frequency effects. Interestingly, some strong associations in the **Pheno∼Env** models were uninformative for the **Pheno∼Inv+Env** models (e.g., PC2_Pheno_ vs. Dim 1_RS_ and among habitat difference). The posterior distributions for these effects were wider (Fig. S8; Fig. S12), which is consistent with a weaker explanatory power in a more complex model.

**Figure 6:**
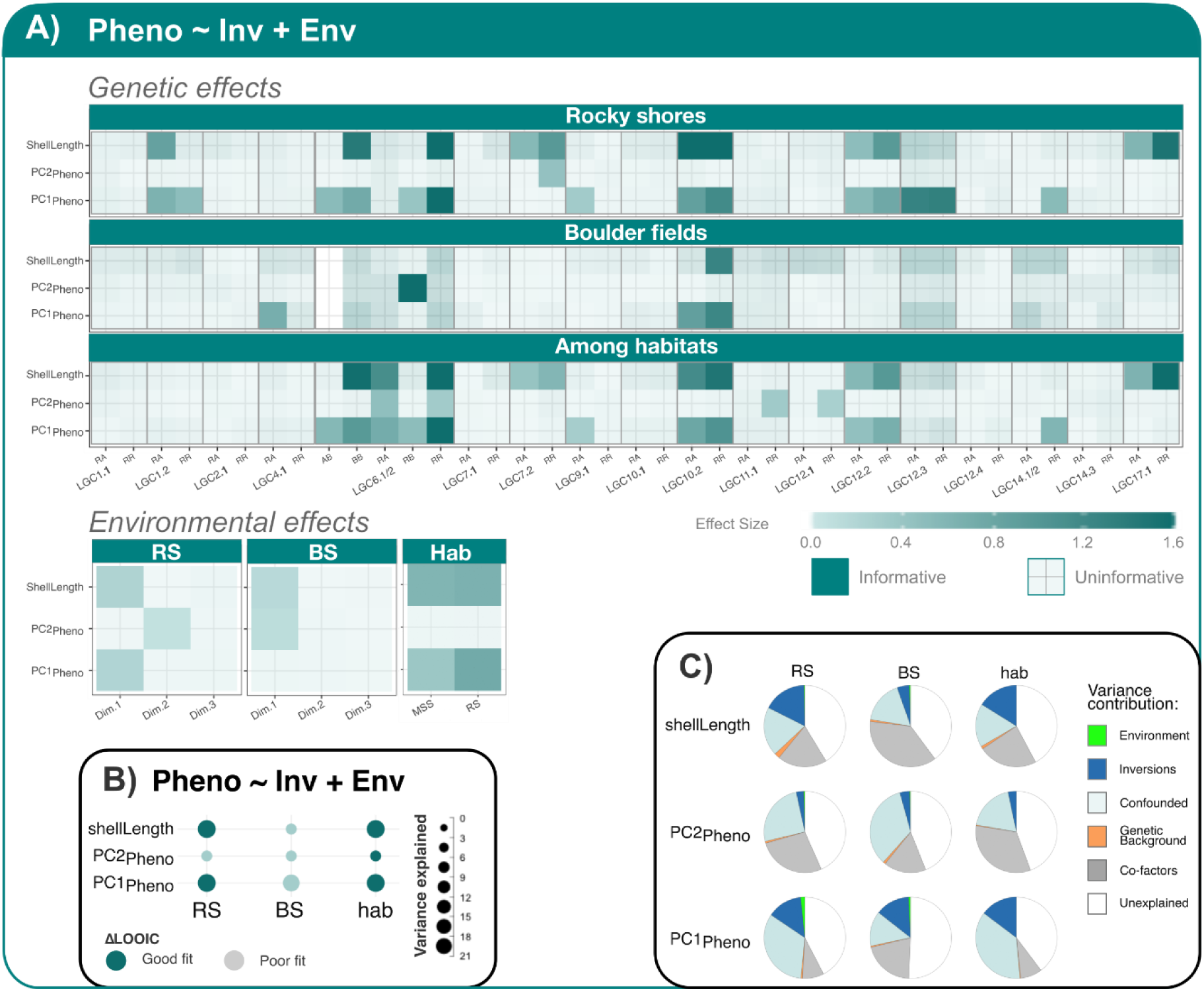
Bayesian regression model results for the **Pheno∼Inv+Env** analysis. **A)** Tile plot summary of **Pheno∼Inv+Env** models. Different habitat results are shown as different sub-panels. Informative parameters were those which did not have credibility intervals (CI) overlapping 0. Posterior distributions are in the supplementary materials (Fig. S12). The effects of each inversion are split into their respective arrangement frequencies relative to the AA arrangement (e.g., RA = β_AA_ – β_RA_; note that there are five possible arrangement contrasts for the complex inversion LGC6.1/2). The effects of environmental PCAmix axes (Dim1-3), and sex and maturity effects, are in separate boxes below **B)** Percentage of variance explained and ΔLOOIC scores for these models. **RS** = rocky shores, **BS** = boulder shores, and **hab** = habitat class models. **C)** Pie chart showing variance decomposition.

Inversion effects on the phenotypes in the **Pheno∼Inv+Env** models were evaluated independently in each habitat type (excluding muddy-sandy shores). LGC12.3 was disregarded due to its low minor arrangement frequency and resulting wide posterior distributions. Five inversions showed no effect on phenotypic variation (LGC1.1, LGC2.1, LGC10.1, LGC12.4 and LGC14.3) in these models despite their association with environmental variation in the **Inv∼Env** models (Fig. 5E, Fig. 6A, Fig. S11). These may represent cases of selection on unmeasured phenotypes (e.g., behaviour, shell colour, physiology or diet preference). Some inversions were associated with a single phenotypic variable; LGC17.1 exclusively influenced shell length, three inversions influenced PC1_Pheno_ (LGC4.1, LGC9.1 and LGC14.1/2), and two influenced PC2_Pheno_ (LGC11.1 and LGC12.1). Three inversions influenced both shell length and PC1_Pheno_ (LGC1.2, LGC10.2 and LGC12.2), LGC7.2 influenced shell length and PC2_Pheno_, and finally LGC6.1/2 influenced all measured phenotypes.

Generally, there were more inversions associated with phenotypic variation in rocky than boulder shores (Fig. 6A). Six inversions had effects only in rocky shores (LGC1.2, LGC7.2, LGC9.1, LGC12.2, LGC14.1/2 and LGC17.1), while only LGC4.1 specifically influenced phenotypes in boulder shores. This pattern is consistent with an environmental gradient that is exclusive to a single habitat, such as the vertical distribution of snails on rocky shores which was previously shown to influence LGC9.1 arrangement frequencies, albeit weakly in Sweden (Morales et al., 2019). Two inversions had phenotypic effects shared across habitats (LGC6.1/2 and LGC10.2). Another two inversions had effects only in the among-habitat models (LGC11.1, and LGC12.1). Both LGC11.1 and LGC12.1 showed strong dependence on the environment in **Inv∼Env** models within both rocky and boulder shores (Fig. 5E), and in turn mainly influenced PC2_Pheno_ (Fig. 6A). These inversions may represent cases where the differences in averages among habitats are stronger than the effects within a habitat, which would be consistent with divergent selection between contrasting habitats.

Variance decomposition showed that a substantial fraction of phenotypic variance can be attributed specifically to the effects of inversions (Fig. 6C). Their overall contribution may be greater, as a large proportion of phenotypic variance was attributed to the confounded effects of environment, inversions and co-factors. This is mostly expected as inversion arrangement frequencies respond to the environment which in turn will impact adaptive phenotypes. Only a small proportion to total phenotypic variance could be attributed specifically to environmental variables, perhaps because the proxies used did not provide a good reflection of the causative environmental axes. More variance was explained by inversions for PC1_Pheno_ and shell length than PC2_Pheno_, but the total variance explained was similar for the three phenotypic axes (Fig. 6C). Only a small proportion of variance in PC1_Pheno_ was exclusively explained by environmental variables, with much potential for environmental effects confounded with inversions and the co-factors in the models (Fig. 6C). For example, 56% of the variance in shell length was explained by among habitat models, with 16% exclusively explained by inversions, 1% explained by the genetic background, 24% explained by co-factors in the model and 17% confounded among factors. No phenotypic variance could be exclusively attributed to the environment (habitat type) for this model.

## Discussion

Using random sampling we have shown that Swedish *L. saxatilis Crab* and *Wave* ecotypes are at opposite sides of a phenotypic continuum, which is primarily linked to wave exposure. Phenotypic variation was pervasive across the landscape we sampled although snails on rocky shores tended to be more variable than those on boulder and muddy-sandy shores. Beyond the earlier defined ecotypes, populations also show strong patterns of local adaptation to continuous habitat variation, especially on a gradient from wave exposed to wave protected rocky shores. Chromosomal inversions strongly influenced phenotypic variation, in response to environmental variation, suggesting the involvement of all 18 known polymorphic inversions in local adaptation.

### Ecotypes in a landscape

*Littorina saxatilis* ecotypes have been extensively studied at the abrupt boundaries between crab haunted and wave exposed habitats (Faria et al., 2019; Hearn et al., 2022; Janson, 1982; Janson & Sundberg, 1983; Koch et al., 2021; Ravinet et al., 2016; Westram et al., 2018, 2021), but the *Crab* and *Wave* ecotypes make up only a part of this species’ phenotypic diversity (Reid 1996). Some of the snails we collected, which were found in muddy shores, matched the descriptions of the comparatively understudied *Brackish* (“*tenebrosa*”) ecotype (Janson & Ward, 1985; Barnes, 1993; Johannesson & Johannesson, 1996; Reid, 1996). Intermediate forms are typically found in hybrid zones among ecotypes (Larsson et al., 2020; Janson & Sundberg, 1983), but these hybrids only occur over very restricted ranges, forming a small minority of the snails that we have sampled. Other forms of snails appear in sites far from hybrid zones and may be the product of variation within a habitat type. Differences among sites of the same habitat type can result in strong fitness differences, like those observed in the *headland* ecotype of *Senecio lautus* (James et al., 2021b, 2025), and it is likely that this is also occurring between *L. saxatilis* populations. Applying a random sampling design over a heterogenous landscape allowed us to observe this broad gradient of phenotypic variables. At abrupt transitions in this landscape, *L. saxatilis* ecotypes are maintained through strong dispersal-selection balance and genome-wide barriers to gene exchange (Westram et al., 2018, 2021) but, elsewhere, populations form dispersal-independent clines (Barton & Gale, 1993), adapting to gradual changes in exposure and predation pressure. This is conceptually similar to the ‘ring ecotypes’ suggested by Johannesson et al. (2026): although *L. saxatilis* is adapting to environmental variation that is more complex than a simple ring.

Phenotypic plasticity may have contributed to the continuous distribution of phenotypic variation. Plasticity could make discrete genetic differences among ecotypes appear as continuous phenotypic variation. A minor plastic component to shell shape has been demonstrated in *Littorina* (Hollander et al., 2006b). However, substantial plastic responses to environmental variation appear to be unlikely since independent environmental effects were weak or absent in **Pheno∼Inv+Env** models (Fig. 6C). Sex and maturity were included as covariates in our analyses which accounted for shape and size dimorphism and growth effects.

A fixed number of snails were sampled at each site, regardless of the local population density, this sampling strategy may have impacted the distribution of phenotypic variation. Densities appeared to be highest on boulder and rocky shores facing the open sea, and thus intermittently exposed to storm surges. Modifying sampling to account for variation in snail density might result in a more clustered distribution of phenotypes, as densities appeared to be higher in typical ecotype habitats relative to intermediate environments. This remains to be tested.

The connected landscape we have shown in *L. saxatilis* may be specific to the region we sampled. Other *L. saxatilis* populations may have more discrete ecotype boundaries than we found in the Kosterhavets National Park, Sweden. Stronger genetic difference was found between Spanish *Crab* and *Wave* ecotypes than between Swedish ecotypes (Kess & Boulding, 2019; Raffini et al., 2025). This can be the result of a different spatial arrangement of distinct habitats, stronger selection gradients, and longer divergence time (Raffini et al., 2025). Iberian *L. saxatilis* ecotype pairs are closer to examples of discontinuous ecotype divergence seen in some other species (Johannesson et al., 2026). Across *Littorina*, different ecotype pairs represent a gradient of divergence resulting from variation in the intensity of gene flow, strength of selection, effective population sizes, and demographic history (Johannesson et al., 2024).

Studies of ecotype pairs in other species have often overlooked alternative or intermediate habitats to focus on divergence at ecotone boundaries or between distinct habitat patches (Johannesson et al., 2026), leading to the perception of ecotypes as discrete groups. In some species, the evidence supports this perception, such as *Senecio lautus* (James et al., 2021a,b; Roda et al., 2013, 2017), Atlantic cod (*Gadus morhua*; Kristensen et al., 2021; Berg et al., 2017; Henriksson et al., 2026), and *Littorina fabalis* (Le Moan et al., 2024). For example, sharp genetic differences have been found between ecotypes of *L. fabalis* despite apparently shallow environmental gradients in the same boulder shore habitats where we sampled *L. saxatilis* (Le Moan et al., 2024). However, not all ecotype pairs fit the model of discrete genetic groups. Gene flow between ecotypes has been observed in pairs of *Timema cristinae* ecotypes, with intermediate populations acting as bridges (Comeault et al., 2015). Threespine sticklebacks (*Gasterosteus aculeatus*) have aspects of both sharp ecotype differences and gradual ongoing gene flow, with these two scenarios playing out among three well-studied ecotype pairs (marine-freshwater, benthic-limnetic and lake-stream; Reid et al., 2021). Some pairs, like the lake-stream ecotypes (Baebler et al.,2026), and the benthic-limnetic ecotypes in lakes (Nagel and Schluter 1998), are maintained despite the close proximity of these habitats, albeit with most lakes having some hybrids (Willacker et al., 2010). Other pairs like marine-freshwater ecotypes are frequently connected through seasonal bouts of gene flow (Schluter & Conte, 2009; Ravinet et al., 2015). Ecotypes are better described on a gradient of divergence, where discrete differences likely indicate presence of intrinsic reproductive barriers formed after earlier periods of geographic isolation (Bierne et al., 2013; Le Moan et al., 2024), or stronger habitat-based selection at abrupt ecotype boundaries (Johannesson et al., 2026). The spatial scale of sampling also modifies signals of local genetic structure (Levin, 1992; Rogivue et al., 2025), and statistically significant environmental associations (Guillaume et al., 2024; Guillaume, 2024), which can blur discrete ecotypes into a gradient at broader spatial scales. Discrete populations at ecotones are thus a null hypothesis which can be tested, rather than assumed as an example of incipient speciation.

### Adaptation across overlapping environmental gradients

In our study area of *L. saxatilis*, other environmental gradients, in addition to the wave exposure gradient, likely have independent adaptive effects. Some of these are captured by the environmental dimensions that we analysed, but others may not be. Shore height is known to contribute to local adaptation in *L. saxatilis* (Morales et al., 2019). High shore snails that live above the waterline, have adapted to survive for longer in warm and dry conditions (Sokolova et al., 2000). Past studies have seen strong phenotypic (Johannesson & Johannesson, 1990) and genetic differences in high shore snails, such as the change in allele frequencies at the allozyme locus *Aat* (Johannesson et al., 1995; Johannesson & Tatarenkov, 1997; Mittermayer et al., 2023). Shore height has therefore been hypothesised as an additional environmental gradient which is perpendicular to the ecotype gradient in Sweden (Westram et al., 2018; 2021; Morales et al., 2019).

Shore height, however, had a limited effect on phenotypes and inversion frequencies in our study. Adding snail height on shore into the **Pheno∼Env** and **Inv∼Env** models for rocky shores had some impact (see supplementary method 4; Fig. S14). However, only the **Inv∼Env** model for the LGC12.2 inversion improved when snail height was added as a term (Table S2). Given that rocky shore transect heights were highly variable in the field, this association was likely driven by only the most exposed sites, which had the greatest shore height ranges in this part of Sweden. The limited effect of shore height more generally may be the result of our random sampling design. Swedish shores are typically flatter (81% of sampling sites had a height range < 1m) than Spanish and British shores. The previously documented adaptive responses to height gradients in Sweden (Johannesson & Johannesson, 1990; Johannesson & Tatarenkov, 1997; Morales et al., 2019; Westram et al., 2018; 2021) were specific to shores representing the more extreme wave exposure.

Unmeasured environmental gradients, other than shore height, could also have influenced inversion arrangement frequencies and phenotypes. For instance, shore substrate, temperature, and season, were found to be informative factors in a species distribution model (Bosso et al., 2022). Our environmental measurements contained some attributes of these gradients (e.g., sampling date roughly controlled for seasonal changes), however continent-spanning gradients used by Bosso et al. (2022) may only loosely correlate with the regional level environmental gradients we sampled. Interactions with other species are likely to contribute to these unknown gradients, for example predator density (European shore crab; *Carcinus maenas*) changes among habitats (supplementary methods 7; Fig. S13), with known effects on the nominal *Crab* ecotype (Johannesson, 1986; Boulding et al., 2017). Analysing these alternative adaptive gradients within a single ecotype can be informative, as this approach may help to break apart the correlations among variables which shift together across an ecotone. Our random sampling approach partially achieved this objective but more detailed tests, with a finer scale of sampling and more detailed characterization of the habitat could connect our observations to specific adaptive environmental gradients.

### The role of inversions in a heterogenous landscape of local adaptation

Arrangement frequencies for multiple chromosomal inversions in *L. saxatilis* change steeply across previously studied hybrid zones (Westram et al. 2018, 2021; Raffini et al. 2025). Coupling makes it difficult to dissect the drivers of individual clines across hybrid zones, although associations with adaptive phenotypes support a role for divergent, habitat-based selection for some inversions. Here, our random sampling spans continuous phenotypic and environmental variation, providing an opportunity to separate out the roles of individual inversions. Overall, inversions were broadly associated with every phenotypic and environmental variable measured. This includes three inversions (LGC6.1/2, LGC14.1/2, and LGC17.1) that contain quantitative trait loci (Koch et al., 2021, 2022), and have previously been associated with ecotype divergence (Morales et al., 2019; Raffini et al., 2025; Reeve et al., 2024; Westram et al., 2021). Phenotypic variation was also associated with over half of the inversions, with the strongest effects from LGC6.1/2. This included inversions that have not been associated with phenotypic variables in previous analyses, such as LGC7.2 (Fig. 5B). However, not all phenotype-inversion effects were included in our study. For example, QLT analysis showed a genetic basis for boldness in *L. saxatilis* (Koch et al., 2021, 2022), a behaviour which is presumably linked to the absence of crab predation pressure in the *Wave* habitat (Johannesson 1986; Johannesson & Johannesson, 1996). Unmeasured physiological and behavioural phenotypes may be influenced by inversions; further study could find that inversions have even wider adaptive roles than we have presented here.

Variable patterns of adaptation were observed when individual inversions were compared (Fig. 6A). Some inversions had general effects across different habitat types (e.g., LGC6.1/2) while others were only associated with local adaptation within a single habitat (e.g. LGC4.1 was involved in adaptation to wave exposure in boulder shores). There was also a third category of inversions where the variation within a single habitat likely contributed to the differences among habitats. For example, mean arrangement frequencies of LGC17.1 were different among rocky shores and boulder shores, with this difference driven primarily by extremely wave-exposed rocky shores. Only two inversions showed differences in arrangement frequencies among but not within habitat types (LGC11.1, and LGC12.1). Somewhat analogous results have been seen in *Helianthus* sunflowers, where inversion polymorphisms that vary in frequency along a latitudinal gradient in *H. annuus* are also associated with ecotype divergence between discrete habitat patches in *H. petiolaris* (Todesco et al., 2020). However, the habitat associations were similar for all inversions, and *Helianthus* hybrid-zones span a much wider geographic scale than in *L. saxatilis*. The prevalence of within habitat effects for *L. saxatilis,* suggests that not only are the effects of inversions pervasive, but that these effects can be decoupled from ecotones and detected via random sampling of habitats. This decoupling indicates that inversions are directly contributing to local adaptation.

The patterns of phenotype-inversion-environment associations we have presented here are possibly an underestimation of the full map of adaptive effects present in this region of Sweden. Multivariate models of heterogenous landscapes tend to underestimate genotype-environment association when loci have pleiotropic effects on phenotypic traits (Lotterhos, 2023), a situation that is presumably common for inversions. Overall, *L. saxatilis* is clearly adapting to more than the *Crab-Wave* ecotype gradient, suggesting that local adaptation is both widespread and responding to diverse environmental drivers. Inversions are major contributors to this heterogenous pattern of local adaptation, with each inversion influencing multiple phenotypes and responding to multiple environmental factors, which is consistent with inversions maintaining large tracks of linkage disequilibrium among a myriad of adaptive loci (Kirkpatrick & Barton, 2006; Berdan et al., 2023).

### Variance decomposition: the role of inversions after considering other factors

Variance decomposition of **Pheno∼Inv+Env** models showed that inversions contributed more to the total explained phenotypic variance than environmentally induced variation. The largest component of phenotypic variance was unexplained by our measured variables. Some unexplained variance may be measurement error, especially given the limitations of ShellShaper when modelling the aperture (Jenny Larsson, personal communication). Another source of unexplained variance comes from loci outside inversions not included in the “genetic background” which may have large adaptive effects (Westram et al., 2018, 2021; Koch et al., 2022); like the situation in threespine sticklebacks where much of the ecotype-associated genetic variance occurs in large effect loci outside of inversions (Peichel & Marques, 2017). In fact, Koch et al. (2022) found loci outside inversions in *L. saxatilis*, which collectively have as large an effect as inversions on phenotypic variation over *Crab*-*Wave* hybrid zones. Currently the adaptive role of specific ‘colinear outliers’ has yet to be investigated further than observations of allele frequency differences (Westram et al., 2018, 2021; Koch et al., 2022).

Much of the explained phenotypic variance was confounded among factors, or exclusively attributed to co-factors (sex, maturity, sampling date; Fig. 6C). Sexual dimorphism has been reported before, caused by sexual selection arising from males tending to mate more successfully with females slightly larger than themselves (Perini et al., 2020). Shells also change shape with age, as a consequence of logarithmic spiral growth (Larsson et al., 2020), with a common garden experiment suggesting that shell development is different between ecotypes (Hollander et al., 2006b). Results of the **Pheno∼Inv+Env** models showed that maturity had a stronger effect on shell size and shape than sex (Fig. S11), indicating that developmental changes may be one of the strongest factors influencing shell shape. After accounting for unexplained variance, confounds and co-factors, the largest remaining source of phenotypic variance was the inversions.

## Conclusion

Ecotypes in Swedish *L. saxatilis* are part of a landscape of local adaptation comprising both sharp environmental breaks and continuous gradients, that fits a conceptual model somewhere between a Harrisonian mosaic (Harrison & Rand, 1989; Ross & Harrison, 2002) and ring species or ring ecotypes (Irwin et al., 2001, 2005, Johannesson et al., 2026). Abrupt habitat transitions lead to strong local adaptation, producing sharp changes in allele frequencies across many loci (Koch et al., 2021, 2022; Morales et al., 2019; Reeve et al., 2024; Westram et al., 2021; Raffini et al., 2025). Such habitat transitions are recognized as *Crab-Wave* hybrid zones, and hybrid zones between *Crab* and *Brackish* ecotypes (the latter rarely investigated but see Janson & Ward, 1985). Other parts of the landscape are shores with much shallower selection gradients essentially making up a single habitat class. Here, alternative phenotypes, either similar to the three main ecotypes or some intermediate phenotype, are dominant (Johannesson et al., 2024; 2026). We found that 12 chromosomal inversions contribute to phenotypic variance, and all 18 inversions have some association with local adaptation in *L. saxatilis*, in the face of pervasive gene flow across a heterogenous environment. By leveraging random sampling, we have mapped out the widespread associations between genotype, phenotype and environmental variation across this landscape. Exploring ecotype pairs in other species using a mix of random sampling and targeted ecotype contrasts will help to contextualise whether *L. saxatilis* ecotypes are exceptional or exemplify a general pattern of ecotype formation.

## Data availability

Raw sequencing data has been uploaded to the NCBI (PRJEB51226). All scripts for processing and analysing data have been uploaded to GitHub (https://github.com/ja-Reeve/Koster_data_analysis).

## Author contributions

JR, KJ and RKB conceived this project. JR and KJ collected snails. JR and OOM extracted DNA and tissue for sequencing. EK and RB designed the SNP panel for genotyping inversions. PB conducted the GiS analysis. JR ran the Bayesian analysis and wrote this manuscript under the guidance of RKB and KJ. All authors commented on the final version of this manuscript.

## Acknowledgments

We would like to thank the *Littorina* Research Community and Centre for Marine Evolutionary Biology (CeMEB) for their advice and feedback for this project. Additionally, we thank Mayram Hoseynzadeh for her help with snail dissections and DNA extractions. Jenny Larsson provided great advice for analysing shell photos. Jon Havenhand, provided practical advice on Bayesian regression modelling. Maddie James gave her expert review to an early version of the discussion. Finally, thank you to Cynthia Riginos and her lab at the University of Queensland, for providing a space for an itinerant scholar to share ideas and learn.

