## Supplementary materials for "The role of chromosomal inversions in local adaptation and ecotype formation across a variable landscape in an intertidal snail"

### Supplementary method 1: drone photo measurements of environments

Each sampling site was photographed by flying a SwellPro SPRY+ drone 10, 20 and 50m above the snail sampling transect. These drone photos were analysed for a range of measurements (Main text Table 1) in ImageJ (Schneider et al., 2012). Not all measurements could be made at each site, so sites were split into three habitat classes: rocky shores (RS), boulder shores (BS) and muddy-sandy shores (MSS). Here each of the measurements will be described. The habitat classes that each measurement applies to are in square brackets next to the name.

**HTD [RS, BS, MSS]:** habitat transition distance was measured from the bottom of the snail sampling transect to the closest habitat transition (e.g., rock to boulder, boulder to sand). Distances were measured in meters following the contours of the shoreline in 1m steps. Any transects that ran across a habitat transition were recorded as 0m. Maximum distance was limited to 50m.

**LVD [RS, BS, MSS]:** land vegetation distance was used as an indicator of wave and wind exposure. Longer distances were expected for more exposed sites. Distance was recorded in meters by drawing a line from the edge of the seaweed bed (or barnacle belt at very exposed sites) to the start of the nearest large patch of land vegetation. These lines were drawn perpendicular to the shoreline starting within 1m of the bottom of the snail transect. Five lines were drawn at each site using, their average as the final measurement. Maximum distance was limited to 50m. Sites without land vegetation (e.g., small skerries) were set to 50m.

**BZW [RS, BS]:** black zone distance was another indicator of wave and wind exposure. It was measured as a subset of the LVD lines, going from the edge of the seaweed bed or barnacle belt to where the black littoral biofilm ends. Sometimes this 'black zone' was patchy. In these cases, the distance to the furthest visible patch of black biofilm was used. This was not recorded in muddy-sandy shores, as the black zone was either absent or difficult to identify without extensive stretches of rocky substrate.

**TZW [MSS]:** tide zone distance was an equivalent measure to BZW for muddy-sandy shores. The end of the tidal zone was the damper area of sand or mud, which marks the upper portion of the foreshore.

**FW [RS, BS, MSS]:** *Fucus* belt width was an indicator of wave exposure with a broader belt of fucoids in moderately exposed shores, absence under strong wave exposure, and patchy or absent in lagoons. The width of the *Fucus* sp./*Ascophyllum* algae belt was measured perpendicular to the shoreline. For complex shorelines, the lines were measured 180° opposite to the direction of waves indicated by the black zone. Five measurements were made within 1m of the bottom of the snail transect. In patchy environments the width was measured across gaps of up to 1m in the *Fucus* belt. The final measurement was the mean. All measurements were in meters. Sites without *Fucus* were scored as 0m.

**FP [RS, BS, MSS]:** *Fucus* belt patchiness was a rough proportion of seaweed present out of the overall area of the *Fucus* belt in each drone photo. Intact belts with FP = 1.0 were indications of moderately exposed sites.

**RF [RS]:** rock fissures were counted along three 5m lines which ran parallel to the middle of the snail transect (i.e., between points D and E). ImageJ's multipoint tool was used to count each fissure under a line. The rounded mean of the three lines was used as the final measurement.

**GS [BS]:** grain size of boulders was an indication of how large boulders were in boulder shores. Like **RF**, three lines were drawn perpendicular to the snail sampling transect, and the number of gaps between boulders was counted. Sites with larger boulders had fewer gaps. In sites with areas of rock, the lines were moved to avoid counting the rock. The final score was the rounded average of the three lines.

### Supplementary method 2: DNA extractions and whole genome data processing

A hybrid approach that used whole genome shotgun (WGS) analysis to develop a SNP panel was used to genotype snails. Individual snails were chosen from 100 separate sites for WGS analysis. Sites were selected to be geographically distant and to cover as many habitats as possible. The largest snail was used at each site, so that replacement tissue would be available in case an extraction failed. DNA was extracted from the foot tissue of each snail using a CTAB-chloroform protocol (Panova et al. 2014). After extractions DNA was cleaned using a Zymo Genomic DNA Clean & Concentrator - 25 kit (Zymo Research). DNA quality was inspected with a Nanodrop 3.0 (Thermo Fisher Scientific) and quantified with a Qubit 3.0 spectrophotometer (Thermo Fisher Scientific). DNA samples were sent to BGI genomics, Hong Kong for library preparation and short read sequencing. Libraries were prepared with BGI genomics standard library preparation. They were then run on a DNBSEQ machine to a target of 5X coverage. One snail failed library preparation, and another was a misclassified juvenile *L. littorea*, leaving 98 samples for downstream analysis.

Sequenced reads were processed into SNPs with a BWA-bcftools pipeline. Reads were mapped onto the *L. saxatilis* reference genome (v1; Westram et al., 2018) with **BWA-MEM** (v0.7.17; Li & Durbin, 2009). Mapped reads were sorted with **samtools** (v1.12; Danecek et al., 2021) and duplicates were removed with **Picard** MarkDuplicates (v2.23.4 Broad Institute, 2018). Variants were called with **bcftools** (v1.12; Danecek et al., 2021), removing indels. Stringent filters were applied to the SNPs and only biallelic sites were kept. SNPs were kept if GQ ≥ 20, QUAL ≥ 30, 5 ≤ DP ≤ double mean depth per sample, ≤ 20% samples were missing and maf ≥ 0.03. Read trimming was not performed, as no systematic errors or adaptor content were identified after inspecting raw reads with **fastqc** (v0.11.9; Andrews, 2018) and **MultiQC** (v1.10; Ewels et al., 2016).

**Supplementary method 3: designing SNP panel for inversion genotyping**

A SNP panel was designed from the WGS data to genotype all samples. SNP panel markers were divided into three sets; i) inversion markers, ii) colinear outliers for *Crab-Wave* divergence (not analysed in this project) and iii) genetic background markers. SNP selection was run as a two-stage process. First 1000 SNPs were genotyped for 99 samples, then the 351 best-mapping SNPs from those successfully genotyped in the first round were used to genotype the remaining 1,389 samples.

Markers in the first round of genotyping were selected differently for the three sets. Inversion markers were drawn from two sources; an existing set of markers from PoolSeq data identified with the "parallelogram" method (Westram et al., 2023), and from the SNPs in our WGS data which were strongly correlated with inversions. Inversion correlations were identified by calculating Pearson's $\rho$ between the genotypes of a given SNP and PC1 scores from a PCA of all SNPs between the boundaries of each inversion (for LG12 Hearn et al., 2022; otherwise Westram et al., 2021). PCA was run with the *prcomp()* function. Only the ten strongest correlating SNPs were kept for further analysis. This process was repeated for PC2 of LCG6.1/2 and LGC14.1/2, as one arrangement separates out on this axis in complex inversions (Faria et al., 2019). Correlations were generally poor for LGC14.3. The three most correlated SNPs were retained, and then seven random SNPs were selected from within the inversion. As a confirmation step, the genotypes of these SNPs were referenced back to the PCA scatterplot to check that the outer clusters were opposing homozygotes (0/0 or 1/1) and that the central cluster contained heterozygotes (0/1). Colinear outliers were drawn directly from a study looking at *Crab-Wave* differences in Sweden (Westram et al., 2018). Any outlier overlapping with the inversion markers was ignored. Finally, a structured random draw was made from the colinear regions of the genome, to fill out the 1000 SNP allotment. Colinear regions were identified by removing all contigs inside inversions and ≤ 2cM outside inversion boundaries. Then one random SNP was selected from each colinear contig. These were then randomly sampled again to select 1000 – N SNPs; where N is the number of inversion markers and colinear outliers. All scripts for this sorting process are on GitHub (https://github.com/ja-Reeve/Koster_data_analysis/SNP_panel_construction).

Additional SNPs were filtered out of the genetic background panel for downstream analysis. 20 SNPs found between 15-47cM on LG5 were removed, as they were inside a putative structural variant (LGC5.1). This structural variant did not impact phenotypic variation but had a minor impact on population structure (Fig. X). Similarly, Contig70288_47522, was also removed after initial testing, as it had a disproportionate effect on a PCA of the genetic background SNPs.

A list of marker positions and snail foot tissue were then sent to LGC Genomics (Berlin, Germany) for DNA extraction and KASP genotyping. The final SNP set was designed from the results of the first round of genotyping. SNPs were filtered to keep those with < 5 missing samples and without a fixed allele. If two or more markers were on the same contig, then one was randomly kept. Inversion markers were filtered to keep those with the tightest correlations, which indicate the strongest linkage disequilibrium with the inversion. Some manual filtering was needed to remove some remaining poorly correlated SNPs, and to sort out the positions of SNPs which were found next to the boundaries of LGC10.1 - 10.2 and LGC14.1/2 – LGC14.3. The final set of SNPs was then sent back to LGC Genomics for KASP genotyping of the remaining samples.

After receiving the SNP genotypes, the final set of SNPs was identified in the original 99 WGS samples, and their genotypes were added to the genotype table. This combined genotype table was further filtered to remove SNPs that were missing in more than 30% of samples. Any remaining missing genotypes were imputed using *knncatimpute()* from the scrime R package (Schwender and Fritsch 2026).

**Supplementary method 4: snail height on shore effects on phenotypes and inversion arrangement frequencies**

Previous studies of Swedish shores show that *L. saxatilis* adapt to fine scale shore height gradients (Morales et al., 2019; Westram et al., 2021). While transect height gives us some information, the position of each snail is also relevant. It was, therefore, important to test how snail height on shore affects phenotypic variation and inversion frequencies.

As an initial test, snail height was added as a co-factor in a subset of brms models. Each snail was given an ordinal rank along the shore height transect at each site. The lowest shore snail was rank 1. Ascending ranks were given to snails until the highest on shore which was rank 8. Snail height ranks were added as a population-level variable to the rocky shore **Pheno~Env** models:

$$Pheno \sim Dim1_{RS} + Dim2_{RS} + Dim3_{RS} + sex + maturity + snail\ height + (1|genetic\ covariance) + (1|Date)$$

Snail height on shore had an effect on phenotypes (Fig. S14) but explained at most 2% of phenotypic variance in all tested models (Table S2). Any adaptive effect of the height gradient seems to have been diluted across sites, as 87% of sampling sites had a height range of < 1m. Previous observations of the height as an adaptive gradient may reflect specific sampling focused on sites with steep rocky shores and larger height ranges. These sites make up only a small part of the Swedish shore habitat of *Littorina saxatilis*. Across the whole region, shore height has only minor effects on shell morphology. However, it is an open question if the shore height is a minor factor in other regions, like Spain, UK or Norway, which have steeper shores and greater tidal ranges.

A similar approach was also used to test for associations between snail height on shore and the two height associated inversions; LGC9.1 and LGC12.2 (Morales et al., 2019; note that Morales et al., 2019 refer to LGC12.2 as LGC12.1. Inversion names on chromosome 12 were updated by Hearn et al., 2022). **Inv~Env** models for rocky shores were modified to add the snail height term, as described above. Like the phenotype tests, shore height had a weak effect on LGC9.1, explaining 1% of the variance among arrangements. However, the effects on LGC12.2 were stronger (Fig. S15), explaining 6% of variance with improved model performances (evaluated as $\Delta$LOOIC; Table S2). Despite this improvement, shore height was not added to the final models in the main analysis because adding height to the LGC12.2 model had little effect on the estimated effects of the other environmental dimensions (e.g., effect of Dim $2_{RS}$ in the **LGC12.2~Env** model = -0.14 [-0.22 – -0.06; Fig. S11], and in the **LGC12.2~Env+height** model = -0.15 [-0.23 - -0.06; Table S2]). Furthermore, the height range was small in the great majority of our sample sites and previous data (Morales et al., 2019; Hearn et al., 2022) did not lead us to expect effects on other inversions.

**Supplementary method 5: variance decomposition**

Phenotypic variance explained by the predictors in the **Pheno~Inv+Env** models was calculated by subtracting factors one at a time in sub-models. Considering each term, and confounded variance among them, would require fitting 277 sub-models per phenotype. We simplified to a handful of models by grouping together factors in four major groups: environmental effects (*Env*), inversion effects (*Inv*), the group-level effects of the genetic background (*gbg*), and the combined effects of the co-factors sex, maturity and the sampling date (*cf*). Each **Pheno~Inv+Env** model thus can be abbreviated as:

$$Pheno \sim Env + Inv + gbg + cf + \varepsilon$$

where Pheno = {$PC1_{Pheno}$, $PC2_{Pheno}$, shellLength}, and ε is the phenotypic variance not explained by the model (i.e., the error term).

Variance explained was estimated for each sub-model using the R package brms. Bayes $R^2$ estimates were made for each draw of a brms model, summarising each model with the average and standard deviation. Bayes $R^2$ from the **Pheno~Inv+Env** models (i.e., full models) was:

$$R^2_{full} = Bayes\ R^2(Pheno \sim Inv + Env + gbg + cf + \varepsilon)$$

The **Pheno~Inv** and **Pheno~Env** models were then used to calculate the variance explained exclusively by environment:

$$R^2_{Env} = R^2_{full} - Bayes\ R^2(Pheno \sim Inv + gbg + cf + \varepsilon) = R^2_{full} - R^2_{Pheno \sim Inv}$$

the variance exclusively explained by inversions:

$$R^2_{Inv} = R^2_{ful} - Bayes\ R^2(Pheno \sim Env + gbg + cf + \varepsilon) = R^2_{full} - R^2_{Pheno \sim Env}$$

the variance exclusively explained by the genetic background:

$$R^2_{gbg} = R^2_{full} - Bayes\ R^2(Pheno \sim Inv + Env + cf + \varepsilon)$$

and finally, the variance exclusively explained by co-factors:

$$R^2_{cf} = R^2_{full} - Bayes\ R^2(Pheno \sim Inv + Env + gbg + \varepsilon)$$

Confounded variance among factors was determined by subtracting the variance explained by each major group of factors from the total variance fitted by the **Pheno~Inv+Env** model:

$$R^2_{conf.} = R^2_{full} - (R^2_{Env} + R^2_{Inv} + R^2_{gbg} + R^2_{cf})$$

The final component was the unexplained phenotypic variance:

$$R^2_{unexp.} = 1 - R^2_{full} = 1 - (R^2_{Env} + R^2_{Inv} + R^2_{gbg} + R^2_{cf} + R^2_{conf.})$$

**Supplementary method 6: comparing environmental variation among habitats**

Anecdotal evidence from the field suggested that rocky shore habitats were more variable than boulder or muddy-sandy shores. Wave exposed sites had less vegetation and greater vertical distances for the littoral zone. Sheltered rocky shores on the other hand, had littoral zones only a few meters wide. To test these observations analytically the variance among sites within each habitat type was compared on a common set of environmental axes. An additional PCAmix analysis was run for all samples using only the variables shared among all three habitats (sin_aspect, cos_aspect, Exposure_FETCH, HTD, LVD, FW, FP, slope, Shore_length, *Fucus*, *Ascophyllum*, Barnacles; Table 1). See the main text for details about the PCAmix approach.

There was more variation within rocky shores than within boulder or muddy-sandy shores. Most of this variance difference was on the first environmental PCAmix dimension (Fig. S5). This dimension was loaded with terms related to wave exposure, thus the greater environmental variation in rocky shores reflects rocky shores occurring across a wider exposure gradient than boulder or muddy shores.

**Supplementary method 7: pilot study to estimate shore crab predation**

Predation from European shore crabs (*Carcinus maenas*) is known to be a major environmental variable driving local adaptation in *L. saxatilis* (Boulding et al., 2017; Johannesson, 1986). The *Crab* ecotype is named for this environmental gradient. In practice measuring crab predation is logistically complicated. Laboratory experiments or tethering in the wild can be done to measure predation, but these experiments are impractical to run across multiple sites. Instead features that are thought to correlate with crab densities are used to estimate predation pressure

(Westram et al., 2018), such as the presence of fucoid seaweeds, which the crabs can use for shelter.

To get a more detailed estimate of crab predation a pilot study was conducted to estimate crab densities near to snail sampling sites. These trials, while quicker to run than tethering experiments, were still found to be too time-consuming to run across all 200 random sampling sites. Due to the small sample size of this pilot study, we decided to remove crab densities from the main analysis. The methods and results of this pilot study are presented here to show the main trend which provides some additional understanding of local adaptation in snails.

Crab densities were surveyed at 18 of the 200 sampling sites. The original sites were divided into six exposure categories based on environmental features (Table S3). Three sampling sites were selected for each exposure class, trying to spread them out across the total sampling region. At each site, three trials were conducted separately by two observers, for a total of six observations. For each trial an open oyster was affixed in the centre of a metal hoop, which was placed at a set distance (0, 10 or 20m) along the shore from the base of the snail sampling transect. After a 5min acclimation period, the observer watched the hoop for 5min counting each crab that entered the area. If a crab was observed re-entering the hoop, it was not counted again. The depth and substrate of oyster placement were also recorded while sampling. Final crab counts were the sum of observations per site.

Crab counts were compared to the first three phenotypic principal components and environmental dimensions of each habitat class (described in the main text), using Spearman’s rank correlation tests.

Crabs were more common in more sheltered shores (Fig. S13A). The total crab count was highest at H6 sites, which were the most sheltered sites. In the other direction, the most exposed sites (H1) never had crabs. These were also rocky shores with a steep drop off and no fucoid cover, which are unsuitable conditions for crabs. Dim $1_{RS}$, shell length and $PC1_{Pheno}$ were significantly correlated with crab counts (Table S4; Fig. S13B-C), which suggests that crab densities predominately vary in rocky shores, which is consistent with our interpretation of $PC1_{Pheno}$ as a Crab-Wave axis and of Dim $1_{RS}$ as an axis related to the exposure-predation gradient. However, crab densities were collected over too few sites to get enough power for this analysis.

The pilot study showed that this was an effective way to measure crab densities, and there was a general trend towards more crabs in sheltered sites. In the field, we also observed more crabs in sheltered areas, but this may be an observation bias. *C. maenas* is very shy, staying under boulders or foliage to avoid capture by birds. We may have underestimated crab densities in sites with thick belts of fucoid seaweed. Expanding the trial to more sites may help to improve our estimates of densities, which may help to explain some of the unexplained phenotypic variance we observed in boulder shores (Fig. 6C).

**Table S1:** Sampling sites descriptions. Latitude and longitude are in decimal degrees (WGS 84). Date is formatted as day-month-year. **N** = sampling count. **Habitat** = habitat type of each sampling location; **RS**= rocky shores, **BS** = boulder shores and **MSS** = muddy-sandy shores.

| Site | Name | Latitude | Longitude | Date | N | Habitat |
|---|---|---|---|---|---|---|
| KT001 | Kattholmen | 58.8658 | 11.1400 | 29-04-2020 | 7 | RS |
| KT002 | Southeast Saltö | 58.867 | 11.13943 | 29-04-2020 | 7 | BS |
| KT003 | North Saltö | 58.87649 | 11.1379 | 29-04-2020 | 7 | RS |
| KT004 | Northwest Saltö | 58.87921 | 11.11954 | 29-04-2020 | 8 | RS |
| KT005 | South Silebukten, Saltö | 58.8778 | 11.12586 | 29-04-2020 | 8 | MSS |
| KT006 | North Silebukten, Saltö | 58.87857 | 11.12791 | 29-04-2020 | 7 | MSS |
| KT007 | North Ängklåvbukten, Saltö | 58.87127 | 11.11775 | 02-05-2020 | 8 | RS |
| KT008 | West Saltö | 58.87258 | 11.11672 | 02-05-2020 | 8 | RS |
| KT009 | Central Ängklåvbukten, Saltö | 58.8706 | 11.12009 | 02-05-2020 | 7 | BS |
| KT010 | Innre Vattenholmen | 58.8767904 | 11.1139289 | 02-05-2020 | 8 | RS |
| KT011 | West Innre Vattenholmen | 58.87683 | 11.11299 | 02-05-2020 | 6 | RS |
| KT012 | Northwest Innre Vattenholmen | 58.87754 | 11.11271 | 02-05-2020 | 8 | RS |
| KT013 | East Yttre_Vattenholmen | 58.875258 | 11.1068681 | 02-05-2020 | 7 | RS |
| KT014 | South Ängklåvbukten, Saltö | 58.86968 | 11.11976 | 06-05-2020 | 5 | RS |
| KT015 | Saltpanbukten, Saltö | 58.87603 | 11.13929 | 06-05-2020 | 8 | RS |
| KT016 | North of Bofors Camping, Saltö | 58.88489 | 11.13937 | 06-05-2020 | 8 | MSS |
| KT017 | Störjrona | 58.85959 | 11.1328 | 07-05-2020 | 8 | RS |
| KT018 | Northeast Lilla Lyngholmen | 58.85548 | 11.14101 | 07-05-2020 | 7 | RS |
| KT019 | Southwest Lilla Lyngnholmen | 58.85449 | 11.13857 | 07-05-2020 | 8 | RS |
| KT020 | South Stora Flatskär | 58.85646 | 11.1396 | 07-05-2020 | 0 | RS |
| KT021 | Southeast Stora Flatskär | 58.8571 | 11.13983 | 07-05-2020 | 8 | BS |
| KT022 | Northwest Yttre Vattenholmen | 58.8757803 | 11.1053751 | 07-05-2020 | 3 | BS |
| KT023 | North Långholmen | 58.88815 | 11.11428 | 19-05-2020 | 5 | RS |
| KT024 | Northwest Långholmen | 58.88742 | 11.11375 | 19-05-2020 | 8 | BS |
| KT025 | Southwest Långholmen | 58.8853 | 11.11623 | 19-05-2020 | 6 | BS |
| KT026 | West Lilla Flatskär | 58.8553 | 11.13635 | 20-05-2020 | 7 | RS |
| KT027 | Southwest Rossö | 58.8367732 | 11.1528383 | 20-05-2020 | 0 | MSS |
| KT028 | Southeast Kackholmen | 58.82832 | 11.15772 | 20-05-2020 | 8 | RS |
| KT029 | Southwest Kackholmen | 58.82618 | 11.14571 | 20-05-2020 | 8 | RS |
| KT030 | Southwest Rossö | 58.82881 | 11.14736 | 20-05-2020 | 8 | BS |
| KT031 | North Ramsö | 58.83823 | 11.07942 | 21-05-2020 | 8 | RS |
| KT032 | East Ulvillarna | 58.82345 | 11.07581 | 21-05-2020 | 8 | RS |
| KT033 | West Ulvillarna | 58.82395 | 11.07221 | 21-05-2020 | 7 | RS |
| KT034 | South Ramsö | 58.82498 | 11.05901 | 21-05-2020 | 7 | RS |
| KT035 | South Långeskär | 58.82745 | 11.04946 | 21-05-2020 | 8 | RS |
| KT036 | East Ramsö | 58.82618 | 11.06804 | 21-05-2020 | 8 | RS |
| KT037 | South Småholmarna | 58.90234 | 11.09665 | 22-05-2020 | 8 | BS |
| KT038 | North Småholmarna | 58.9033 | 11.09661 | 22-05-2020 | 8 | MSS |
| KT039 | East Småholmarna | 58.90308 | 11.09796 | 22-05-2020 | 5 | RS |
| KT040 | South Matkullen | 58.88267 | 11.11718 | 25-05-2020 | 8 | RS |
| KT041 | Between Innre & Yttre Tenskär | 58.88206 | 11.12784 | 25-05-2020 | 8 | RS |
| KT042 | North Klunken | 58.88306 | 11.12751 | 25-05-2020 | 8 | RS |
| KT043 | West Södra Öddö near boating sign | 58.89676 | 11.14417 | 28-05-2020 | 8 | BS |
| KT044 | West Södra Öddö near pier | 58.89799 | 11.14442 | 28-05-2020 | 8 | RS |
| KT045 | North Sämundholmen | 58.89964 | 11.14322 | 28-05-2020 | 7 | MSS |
| KT046 | South Norra Öddö | 58.90058 | 11.13488 | 29-05-2020 | 7 | RS |
| KT047 | Northwest Ramnkrokholmen | 58.90026 | 11.13797 | 29-05-2020 | 8 | MSS |
| KT048 | East Styrsö | 58.90855 | 11.1154712 | 29-05-2020 | 7 | BS |

| | | | | | | |
|---|---|---|---|---|---|---|
| KT049 | West Norra Öddö | 58.9087155 | 11.1175222 | 29-05-2020 | 7 | MSS |
| KT050 | Between Burholmarna | 58.8991388 | 11.1144706 | 29-05-2020 | 8 | MSS |
| KT051 | Southwest Lilleskär | 58.88731 | 11.09935 | 15-06-2020 | 8 | RS |
| KT052 | South Krugglö | 58.8891 | 11.09439 | 15-06-2020 | 7 | RS |
| KT053 | Southeast Grötholmen | 58.89174 | 11.1028 | 15-06-2020 | 8 | RS |
| KT054 | West Yttre Burholmen | 58.89635 | 11.11221 | 15-06-2020 | 8 | BS |
| KT055 | Ramsholmarna | 58.89442 | 11.11565 | 15-06-2020 | 8 | RS |
| KT056 | Southeast Tjälleskär | 58.84502 | 11.03504 | 17-06-2020 | 8 | RS |
| KT057 | West Ramsökalven | 58.83066 | 11.03822 | 17-06-2020 | 7 | BS |
| KT058 | Southwest Ramsholmen | 58.844 | 11.04721 | 17-06-2020 | 8 | RS |
| KT059 | South Ramsholmen | 58.84377 | 11.05081 | 17-06-2020 | 8 | BS |
| KT060 | East Tjälleskär | 58.84666 | 11.03742 | 17-06-2020 | 6 | BS |
| KT061 | Southwest Tjälleskär | 58.8468 | 11.03318 | 17-06-2020 | 7 | BS |
| KT062 | East Ramsholmen | 58.84776 | 11.04888 | 17-06-2020 | 8 | BS |
| KT063 | Northwest Ramsholmen | 58.84949 | 11.05032 | 17-06-2020 | 8 | BS |
| KT064 | Lilla Tjälleskär | 58.85112 | 11.0423 | 17-06-2020 | 7 | BS |
| KT065 | Rundegrå | 58.85286 | 11.04743 | 17-06-2020 | 8 | RS |
| KT066 | Gråaholmarna | 58.85474 | 11.0477 | 17-06-2020 | 7 | BS |
| KT067 | Rock Southeast of Dammstånggrån | 58.85738 | 11.04434 | 17-06-2020 | 7 | RS |
| KT068 | South Lilla Sneholmen | 58.88695 | 11.0646 | 18-06-2020 | 8 | RS |
| KT069 | East Stora Sneholmen | 58.89164 | 11.06426 | 18-06-2020 | 7 | RS |
| KT070 | Northwest Stora Sneholmen | 58.89254 | 11.06115 | 18-06-2020 | 6 | RS |
| KT071 | West Stora Sneholmen | 58.89169 | 11.06168 | 18-06-2020 | 8 | BS |
| KT072 | Southeast Hamneholmen | 58.89185 | 11.05739 | 09-07-2020 | 8 | RS |
| KT073 | Southwest Lindholmen | 58.8848 | 11.12262 | 02-07-2020 | 8 | RS |
| KT074 | West Lindholmen | 58.88573 | 11.12169 | 02-07-2020 | 7 | MSS |
| KT075 | North Killingholmen | 58.8935 | 11.13663 | 02-07-2020 | 8 | RS |
| KT076 | East Killingholmen | 58.89087 | 11.13798 | 02-07-2020 | 8 | MSS |
| KT077 | Island in Lidlerran | 58.88972 | 11.14083 | 02-07-2020 | 8 | BS |
| KT078 | Southwest Hamneholmen | 58.89077 | 11.05087 | 09-07-2020 | 8 | BS |
| KT079 | North Hamneholmen | 58.89622 | 11.05198 | 09-07-2020 | 7 | RS |
| KT080 | Southwest Nordkoster | 58.88452 | 10.98942 | 09-07-2020 | 8 | BS |
| KT081 | Northern Teneskär | 58.88011 | 11.0013 | 09-07-2020 | 8 | RS |
| KT082 | North Rundholmen | 58.86469 | 10.99827 | 10-07-2020 | 8 | BS |
| KT083 | Gula skäret | 58.87238 | 10.99375 | 10-07-2020 | 8 | RS |
| KT084 | West Hällsholmen | 58.87675 | 10.99758 | 10-07-2020 | 8 | BS |
| KT085 | North Hällsholmen | 58.87774 | 10.9964 | 10-07-2020 | 7 | BS |
| KT086 | Ölskär | 58.87228 | 11.00472 | 10-07-2020 | 6 | RS |
| KT087 | West Sydkoster | 58.87542 | 11.00989 | 10-07-2020 | 7 | BS |
| KT088 | Rock near southern Teneskär | 58.86861 | 11.013 | 10-07-2020 | 7 | RS |
| KT089 | Southeast Filjholmen | 58.87738 | 11.06311 | 16-07-2020 | 8 | BS |
| KT090 | Northeast Filjholmen | 58.87945 | 11.06143 | 16-07-2020 | 8 | RS |
| KT091 | West Filjholmen | 58.87858 | 11.05913 | 16-07-2020 | 2 | MSS |
| KT092 | East Stora Brattskär | 58.86079 | 11.0762 | 16-07-2020 | 8 | RS |
| KT093 | West Stora Brattskär | 58.86259 | 11.07198 | 16-07-2020 | 8 | BS |
| KT094 | Northwest Stora Brattskär | 58.86349 | 11.07324 | 16-07-2020 | 8 | RS |
| KT095 | West Lilla Brattskär | 58.86268 | 11.06736 | 16-07-2020 | 8 | BS |
| KT096 | North Vedskär | 58.85825 | 11.07099 | 16-07-2020 | 7 | RS |
| KT097 | West Vedskär | 58.85713 | 11.06906 | 16-07-2020 | 8 | RS |
| KT098 | Southwest Vedskär | 58.85633 | 11.06914 | 16-07-2020 | 8 | BS |
| KT099 | Islet in Kölnebukten | 58.88586 | 11.1345 | 20-07-2020 | 8 | RS |

| | | | | | | |
|---|---|---|---|---|---|---|
| KT100 | Islet in Rävlingsundet | 58.88908 | 11.13352 | 20-07-2020 | 8 | RS |
| KT101 | Dammstånggrån | 58.8576 | 11.04259 | 22-07-2020 | 6 | RS |
| KT102 | Eleven | 58.85584 | 11.04252 | 22-07-2020 | 8 | RS |
| KT103 | Tärnholmen | 58.85467 | 11.03399 | 22-07-2020 | 8 | RS |
| KT104 | Islet south of Tärnholmen | 58.85201 | 11.03209 | 22-07-2020 | 7 | RS |
| KT105 | Islet west of Stenholmen | 58.85361 | 11.02882 | 22-07-2020 | 8 | RS |
| KT106 | Timmerglaset | 58.85207 | 11.02519 | 22-07-2020 | 8 | RS |
| KT107 | West Timmerglaset | 58.85358 | 11.02149 | 23-07-2020 | 8 | RS |
| KT108 | Västra Klaseskären | 58.85902 | 11.02485 | 23-07-2020 | 8 | RS |
| KT109 | Lådderskären | 58.86161 | 11.01379 | 23-07-2020 | 8 | RS |
| KT110 | Middle of Brevik, Sydkoster | 58.86978 | 11.03202 | 23-07-2020 | 8 | MSS |
| KT111 | South Brevik, Sydkoster | 58.86757 | 11.02845 | 23-07-2020 | 8 | MSS |
| KT112 | North Tångundden, Sydkoster | 58.86774 | 11.02616 | 23-07-2020 | 8 | BS |
| KT113 | South Tångudden, Sydkoster | 58.8661 | 11.02591 | 23-07-2020 | 8 | BS |
| KT114 | West Innre Arsklåvet | 58.83453 | 11.13648 | 24-07-2020 | 8 | BS |
| KT115 | North Innre Arsklåvet | 58.83604 | 11.13526 | 24-07-2020 | 8 | RS |
| KT116 | West Matkullen | 58.88308 | 11.11599 | 24-07-2020 | 8 | RS |
| KT117 | Klåvskär | 58.90142 | 11.10859 | 24-07-2020 | 8 | RS |
| KT118 | West Långholmen | 58.90241 | 11.10087 | 24-07-2020 | 8 | RS |
| KT119 | Northwest Långholmen | 58.90448 | 11.10028 | 24-07-2020 | 8 | BS |
| KT120 | Between Styrsö and Elingholmen | 58.90854 | 11.10185 | 24-07-2020 | 7 | RS |
| KT121 | Ålekråkeskär | 58.82461 | 11.02781 | 01-08-2020 | 8 | RS |
| KT122 | Måskär | 58.86044 | 10.98382 | 01-08-2020 | 8 | RS |
| KT123 | Islet West of Jutholmen | 58.87264 | 10.98391 | 01-08-2020 | 8 | RS |
| KT124 | Nypholmen | 58.87628 | 10.98721 | 01-08-2020 | 8 | RS |
| KT125 | Hummerskär | 58.85579 | 10.98123 | 01-08-2020 | 8 | RS |
| KT126 | North Kilesand near houses, Sydkoster | 58.88465 | 11.04181 | 14-08-2020 | 8 | MSS |
| KT127 | North Kilesand, Sydkoster | 58.88291 | 11.04264 | 14-08-2020 | 7 | MSS |
| KT128 | South Hattevik. Sydkoster | 58.88739 | 11.05098 | 14-08-2020 | 6 | RS |
| KT129 | Svartskär | 58.92636 | 11.03502 | 17-08-2020 | 8 | RS |
| KT130 | East Skarvesätet | 58.91407 | 11.00965 | 17-08-2020 | 7 | RS |
| KT131 | Southwest Skarvesätet | 58.91242 | 11.00826 | 17-08-2020 | 8 | RS |
| KT132 | South Husevik, Nordkoster | 58.88842 | 10.98209 | 17-08-2020 | 8 | RS |
| KT133 | Husevik, Nordkoster | 58.88842 | 10.98354 | 17-08-2020 | 8 | BS |
| KT134 | Northeast Mårholmen | 58.88679 | 10.9717 | 17-08-2020 | 7 | RS |
| KT135 | South Mårholmen | 58.8822 | 10.96814 | 17-08-2020 | 8 | RS |
| KT136 | Burholmen | 58.84945 | 10.99243 | 17-08-2020 | 8 | RS |
| KT137 | Gofferholmarna | 58.8379 | 11.13449 | 17-08-2020 | 8 | RS |
| KT138 | Ölbäshskär | 58.84587 | 11.1406 | 17-07-2020 | 8 | RS |
| KT139 | Northwest Lilleskär | 58.88829 | 11.10013 | 18-08-2020 | 8 | RS |
| KT140 | South Kipplingarna | 58.91315 | 11.08973 | 19-08-2020 | 8 | RS |
| KT141 | West Kipplingarna | 58.91327 | 11.08907 | 19-08-2020 | 7 | RS |
| KT142 | Koskär | 58.91492 | 11.10577 | 19-08-2020 | 7 | RS |
| KT143 | Near Eskilbågen, Norra Öddö | 58.9064 | 11.15142 | 19-08-2020 | 6 | RS |
| KT144 | West Stigkilen, Norra Öddö | 58.9109 | 11.13625 | 19-08-2020 | 8 | MSS |
| KT145 | East Stigkilen, Norra Öddö | 58.91215 | 11.13953 | 19-08-2020 | 7 | MSS |
| KT146 | East Ramsökalven | 58.83101 | 11.04365 | 25-08-2020 | 6 | RS |
| KT147 | Northeast Ramsökalven | 58.83368 | 11.04557 | 25-08-2020 | 8 | RS |
| KT148 | Stora Skorna | 58.84377 | 11.01292 | 25-08-2020 | 8 | RS |
| KT149 | Hättorna | 58.84922 | 10.99815 | 25-08-2020 | 8 | BS |
| KT150 | Otterstenarna | 58.85557 | 10.99794 | 25-08-2020 | 8 | BS |
| KT151 | Islet near Runnholmen | 58.86242 | 10.99965 | 25-08-2020 | 7 | RS |

| | | | | | | |
|---|---|---|---|---|---|---|
| KT152 | Bockholmen | 58.86451 | 11.00321 | 25-08-2020 | 8 | RS |
| KT153 | South of Lila Sockna | 58.8387 | 11.00991 | 27-08-2020 | 8 | RS |
| KT154 | Skateskären | 58.83912 | 11.00462 | 27-08-2020 | 8 | RS |
| KT155 | Northwest Arholmen | 58.87035 | 10.99898 | 27-08-2020 | 8 | BS |
| KT156 | Långevik, Sydkoster | 58.88449 | 11.00388 | 27-08-2020 | 7 | RS |
| KT157 | Hattevik, Sydkoster | 58.88906 | 11.04875 | 01-09-2020 | 8 | RS |
| KT158 | Central Kilesand, Sydkoster | 58.8788916 | 11.0475723 | 01-09-2020 | 0 | MSS |
| KT159 | East Kyrkosund, Sydkoster | 58.8700 | 11.05811 | 01-09-2020 | 8 | MSS |
| KT160 | West Kyrkosund, Sydkoster | 58.86445 | 11.05246 | 01-09-2020 | 8 | RS |
| KT161 | Prästundden, Saltholmen | 58.863 | 11.04756 | 01-09-2020 | 8 | BS |
| KT162 | Rock near Saltholmen | 58.90955 | 11.02291 | 07-09-2020 | 7 | RS |
| KT163 | Flate-Guri | 58.91069 | 11.0233 | 07-09-2020 | 7 | RS |
| KT164 | North Saltholmen | 58.91493 | 11.01443 | 07-09-2020 | 8 | RS |
| KT165 | Southeast Skarvesätet | 58.9132 | 11.00955 | 07-09-2020 | 8 | BS |
| KT166 | Lökepottan, Nordkoster | 58.9122 | 11.00486 | 07-09-2020 | 8 | RS |
| KT167 | North Hasslevikarna, Nordkoster | 58.91128 | 11.0026 | 07-09-2020 | 8 | RS |
| KT168 | Märebukterna, Nordkoster | 58.91157 | 10.99736 | 07-09-2020 | 8 | RS |
| KT169 | East Ekenäs, Sydkoster | 58.89807 | 11.04274 | 15-09-2020 | 8 | RS |
| KT170 | Islet in Rörvik | 58.89948 | 11.04041 | 15-09-2020 | 6 | RS |
| KT171 | Mouth of Kostersundet | 58.89895 | 11.03017 | 15-09-2020 | 5 | BS |
| KT172 | Brevik harbour, Sydkoster | 58.87197 | 11.02504 | 15-09-2020 | 8 | BS |
| KT173 | Rönnevik, Sydkoster | 58.86464 | 11.04052 | 15-09-2020 | 8 | MSS |
| KT174 | Skabotången, Norra Öddö | 58.92122 | 11.1405 | 15-09-2020 | 8 | RS |
| KT175 | Halserberget, Norra Öddö | 58.92521 | 11.12917 | 15-09-2020 | 8 | BS |
| KT176 | North Lindholmen | 58.89174 | 11.12642 | 17-09-2020 | 8 | MSS |
| KT177 | Furholmarna | 58.89424 | 11.12214 | 17-09-2020 | 8 | RS |
| KT178 | Southeast Yttre Burholmen | 58.8968 | 11.12021 | 17-09-2020 | 5 | RS |
| KT179 | Between Burholmarna | 58.89904 | 11.11631 | 17-09-2020 | 8 | MSS |
| KT180 | Southeast Innre Burholmen | 58.8983 | 11.12527 | 17-09-2020 | 7 | BS |
| KT181 | Krugglö | 58.89006 | 11.09494 | 25-09-2020 | 5 | RS |
| KT182 | East Tjärnö | 58.89264 | 11.14908 | 25-09-2020 | 4 | RS |
| KT183 | Northeast Tjärnö | 58.89445 | 11.14356 | 25-09-2020 | 0 | RS |
| KT184 | East Ramnekroken | 58.90052 | 11.14639 | 27-09-2020 | 6 | BS |
| KT185 | West Ramnekroken | 58.9023 | 11.14465 | 27-09-2020 | 7 | RS |
| KT186 | Middle of Ramnekroken | 58.90292 | 11.14836 | 27-09-2020 | 7 | BS |
| KT187 | Halserberget near Svaneskär, Norra Öddö | 58.92559 | 11.1258 | 30-09-2020 | 6 | BS |
| KT188 | Elingholmen | 58.90807 | 11.10079 | 30-09-2020 | 6 | RS |
| KT189 | Inrre Burholmen | 58.90198 | 11.11893 | 08-10-2020 | 6 | BS |
| KT190 | Långestrand, Norra Öddö | 58.90272 | 11.1242 | 08-10-2020 | 7 | MSS |
| KT191 | Liggstill | 58.89843 | 11.13116 | 08-10-2020 | 7 | RS |
| KT192 | Trollholmen | 58.8967 | 11.14001 | 08-10-2020 | 7 | RS |
| KT193 | West Styrsö | 58.9068 | 11.10573 | 11-10-2020 | 8 | BS |
| KT194 | Southwest Styrsö | 58.90412 | 11.10553 | 11-10-2020 | 6 | RS |
| KT195 | Northeast Långholmen | 58.8880294 | 11.1158088 | 11-10-2020 | 8 | RS |
| KT196 | Saltö-Lyngnholmen | 58.8624298 | 11.1312154 | 11-10-2020 | 8 | RS |
| KT197 | Svinnäs, Lindholmen | 58.88399 | 11.13719 | 11-10-2020 | 8 | MSS |
| KT198 | Rävlingsundet | 58.89037 | 11.13444 | 12-10-2020 | 8 | MSS |
| KT199 | Tidal flat between Tjärnö and Killingholmen | 58.89005 | 11.1379 | 12-10-2020 | 8 | MSS |
| KT200 | Bofors Camping, Tjärnö | 58.88262 | 11.14268 | 12-10-2020 | 5 | MSS |

**Table S2:** Bayesian regression estimates from three models of snail height on shore effects on phenotypes. The snail height models were built upon the **Pheno~Env** models, and the **Inv~Env** models for LGC9.1 and LGC12.2, for rocky shores with the addition of the predictor snail height; a rank of snail positions from 1 at low shore to 8 at high shore. Models were compared with LOOIC ($\Delta$LOOIC = $LOOIC_{model} - LOOIC_{model+height}$) and Bayes' $R^2$ to assess the explanatory power and variance explained by snail height. **Estimate** = Bayesian estimate of parameters. **SE** = standard error of parameter estimates. **2.5%_CI** and **97.5%_CI** = credibility intervals of the posterior distributions. **Informative** = indication of how informative the estimated parameter is to the tested phenotype (Y|N), determined by the credibility intervals overlap 0. See Fig. S14 for posterior distributions. Intercept and group-level parameters (i.e., random factors) are omitted.

| Phenotype | Predictor | Estimate | SE | 2.5%_CI | 97.5%_CI | Informative |
|---|---|---|---|---|---|---|
| $PC1_{Pheno}$ | sex | -0.42 | 0.10 | -0.61 | -0.23 | Y |
| | maturity | -0.59 | 0.11 | -0.80 | -0.37 | Y |
| | $Dim1_{RS}$ | -0.40 | 0.03 | -0.47 | -0.33 | Y |
| | $Dim2_{RS}$ | 0.05 | 0.04 | -0.02 | 0.13 | N |
| | $Dim3_{RS}$ | -0.22 | 0.04 | -0.30 | -0.13 | Y |
| | Snail height | 0.12 | 0.02 | 0.08 | 0.16 | Y |
| | **$LOOIC_{Pheno\sim Env}$:** | 3081.0 ± 46.9 | **$LOOIC_{height}$:** | 3048.3 ± 46.0 | **ΔLooic:** | 16.4 ± 29.6 |
| | **$R^2_{Pheno\sim Env}$:** | 0.44 ± 0.02 | **$R^2_{height}$:** | 0.46 ± 0.02 | **$\Delta R^2$:** | 0.02 |
| $PC2_{Pheno}$ | sex | 0.11 | 0.07 | -0.02 | 0.25 | N |
| | maturity | 0.17 | 0.08 | 0.02 | 0.32 | Y |
| | $Dim1_{RS}$ | -0.08 | 0.03 | -0.13 | -0.03 | Y |
| | $Dim2_{RS}$ | -0.03 | 0.03 | -0.09 | 0.02 | N |
| | $Dim3_{RS}$ | -0.09 | 0.03 | -0.15 | -0.03 | Y |
| | Snail height | -0.00 | 0.01 | -0.03 | 0.02 | N |
| | **$LOOIC_{Pheno\sim Env}$:** | 2434.8 ± 41.3 | **$LOOIC_{height}$:** | 2436.9 ± 41.3 | **ΔLooic:** | -1.1 ± 25.3 |
| | **$R^2_{Pheno\sim Env}$:** | 0.54 ± 0.02 | **$R^2_{height}$:** | 0.54 ± 0.02 | **$\Delta R^2$:** | $2.44\times10^{-4}$ |
| shellLength | sex | -0.93 | 0.16 | -1.25 | -0.61 | Y |
| | maturity | -2.54 | 0.18 | -2.90 | -2.18 | Y |
| | $Dim1_{RS}$ | -0.57 | 0.06 | -0.68 | -0.46 | Y |
| | $Dim2_{RS}$ | -0.07 | 0.06 | -0.20 | 0.06 | N |
| | $Dim3_{RS}$ | -0.28 | 0.07 | -0.41 | -0.14 | Y |
| | Snail height | 0.16 | 0.03 | 0.10 | 0.23 | Y |
| | **$LOOIC_{Pheno\sim Env}$:** | 3978.7 ± 49.4 | **$LOOIC_{height}$:** | 3955.2 ± 47.9 | **ΔLooic:** | 11.7 ± 28.9 |
| | **$R^2_{Pheno\sim Env}$:** | 0.41 ± 0.02 | **$R^2_{height}$:** | 0.43 ± 0.02 | **$\Delta R^2$:** | 0.02 |
| LGC9.1 | sex | -0.02 | 0.12 | -0.26 | 0.21 | N |
| | maturity | -0.02 | 0.14 | -0.30 | 0.25 | N |
| | $Dim1_{RS}$ | 0.21 | 0.04 | 0.14 | 0.29 | Y |
| | $Dim2_{RS}$ | -0.18 | 0.05 | -0.28 | -0.08 | Y |
| | $Dim3_{RS}$ | 0.24 | 0.05 | 0.13 | 0.35 | Y |
| | Snail height | 0.08 | 0.03 | 0.03 | 0.13 | Y |
| | **$LOOIC_{Inv\sim Env}$:** | 1747.3 ± 24.0 | **$LOOIC_{height}$:** | 1513.9 ± 36.5 | **ΔLooic:** | 116.7 ± 19.4 |
| | **$R^2_{Inv\sim Env}$:** | 0.19 ± 0.02 | **$R^2_{height}$:** | 0.20 ± 0.02 | **$\Delta R^2$:** | 0.01 |
| LGC12.2 | sex | -0.56 | **0.11** | -0.78 | **-0.35** | Y |
| | maturity | 0.34 | 0.12 | 0.10 | 0.59 | Y |
| | $Dim1_{RS}$ | 0.07 | 0.03 | 0.00 | 0.14 | N |
| | $Dim2_{RS}$ | -0.15 | 0.04 | -0.23 | -0.06 | Y |
| | $Dim3_{RS}$ | 0.15 | 0.05 | 0.06 | 0.25 | Y |
| | Snail height | 0.18 | 0.02 | 0.13 | 0.22 | Y |
| | **$LOOIC_{Inv\sim Env}$:** | 1747.5 ± 24.0 | **$LOOIC_{height}$:** | 1687.2 ± 24.9 | **ΔLooic:** | 30.2 ± 7.7 |
| | **$R^2_{Inv\sim Env}$:** | 0.16 ± 0.03 | **$R^2_{height}$:** | 0.22 ± 0.03 | **$\Delta R^2$:** | 0.06 |

**Table S3.** crab sampling site details. Wave exposure was assessed using diagnostic environmental features.

| Category | Wave Exposure | Environmental features | Site ID | Latitude | Longitude | Habitat |
|---|---|---|---|---|---|---|
| H1 | Highly exposed | No fucoid seaweeds; barnacle present | KT034 | 58.8250 | 11.0590 | RS |
| | | | KT051 | 58.8873 | 11.0994 | RS |
| | | | KT122 | 58.8604 | 10.9838 | RS |
| H2 | Moderately exposed | *Fucus vesiculosus* without floating bladders | KT019 | 58.8545 | 11.1386 | RS |
| | | | KT124 | 58.8763 | 10.9872 | RS |
| H3 | Semi-sheltered | *Fucus vesiculosus* with floating bladders but no *Ascophyllum nodosum* | KT012 | 58.8775 | 11.1127 | RS |
| | | | KT037 | 58.9023 | 11.0967 | BS |
| | | | KT062 | 58.8478 | 11.0489 | BS |
| | | | KT073 | 58.8848 | 11.1226 | RS |
| H4 | Sheltered | *Fucus vesiculosus* with floating bladders and *Ascophyllum nodosum* | KT043 | 58.8968 | 11.1442 | BS |
| | | | KT087 | 58.8754 | 11.0099 | BS |
| | | | KT101 | 58.8576 | 11.0426 | RS |
| H5 | Highly sheltered | Mud flat without fucoid seaweeds and grass on shore | KT005 | 58.8778 | 11.1257 | MSS |
| | | | KT076 | 58.8909 | 11.1380 | MSS |
| | | | KT126 | 58.8847 | 11.0418 | MSS |
| H6 | Sandy shore | Small patches of fucoid seaweed, often with low *L. saxatilis* densities | KT002 | 58.8670 | 11.1394 | BS |
| | | | KT027 | 58.8368 | 11.1528 | MSS |
| | | | KT091 | 58.8786 | 11.0591 | MSS |

**Table S4:** Spearman's rank correlation test results between phenotypic variables and crab densities, and between environmental dimensions and crab densities.

| Variable | Rho | S | P-value |
|---|---|---|---|
| *Phenotype* (N = 17) | | | |
| $PC1_{Pheno}$ | 0.72 | 231.24 | $1.21 \times 10^{-3}$ |
| $PC2_{Pheno}$ | 0.43 | 462.52 | 0.08 |
| shellLength | 0.58 | 344.35 | 0.02 |
| *Rocky shores* (N = 8) | | | |
| Dim $1_{RS}$ | -0.89 | 159.09 | $2.75 \times 10^{-3}$ |
| Dim $2_{RS}$ | -0.22 | 102.24 | 0.61 |
| Dim $3_{RS}$ | -0.65 | 138.71 | 0.08 |
| *Boulder shores* (N = 5) | | | |
| Dim $1_{BS}$ | 0.20 | 16 | 0.78 |
| Dim $2_{BS}$ | 0.40 | 12 | 0.52 |
| Dim $3_{BS}$ | -0.60 | 32 | 0.35 |
| *Mud-sandy shores* (N = 4) | | | |
| Dim $1_{MSS}$ | 0.60 | 4 | 0.42 |
| Dim $2_{MSS}$ | -0.80 | 18 | 0.33 |
| Dim $3_{MSS}$ | 0.04 | 6 | 0.75 |

**Supplementary Figures:**

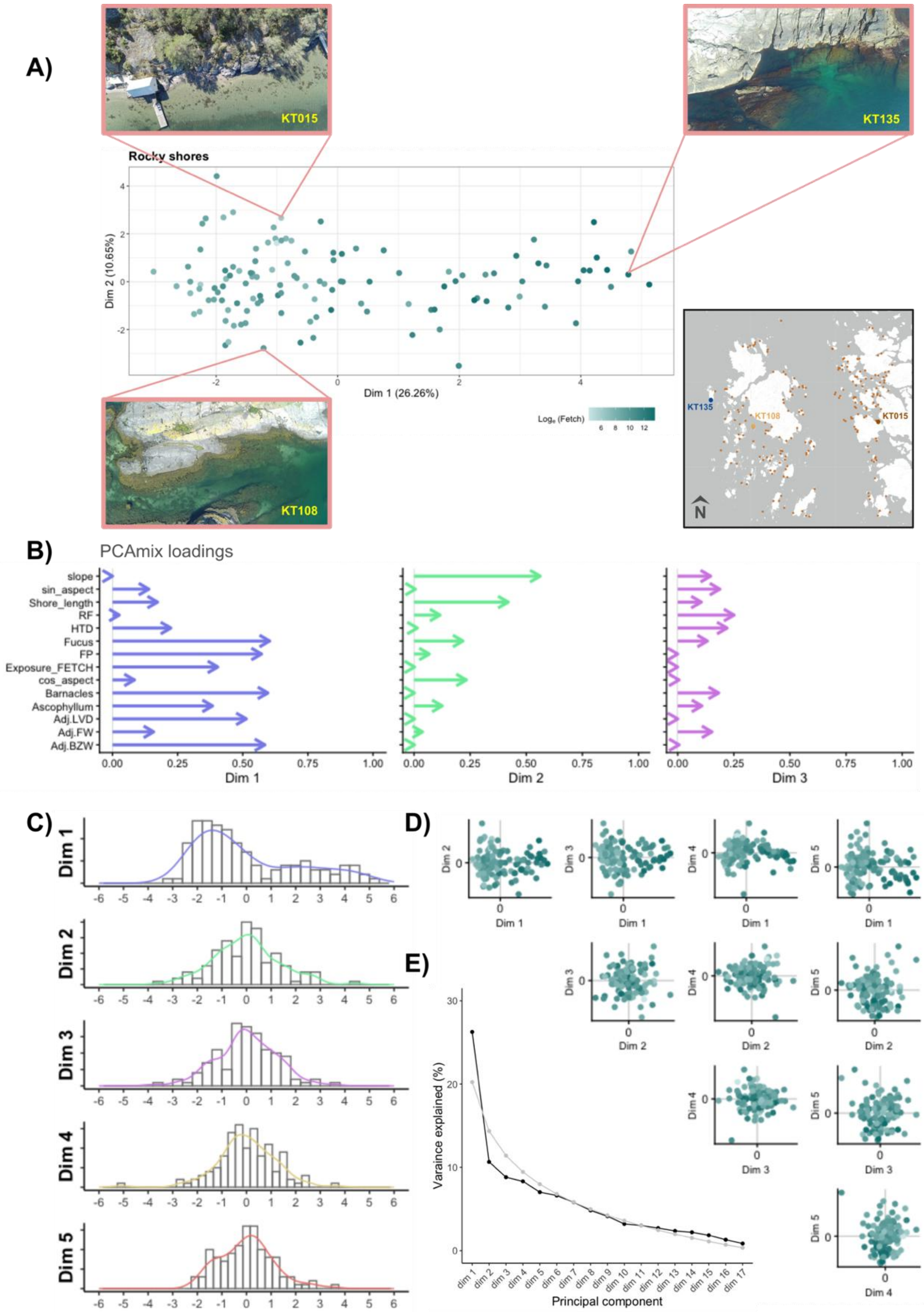


**Figure S1:** PCAmix results for all rocky shores. **A)** Scatterplot comparing PCAmix dimensions 1 and 2. Points are coloured by exposure categories observed in the field. Three drone figures are shown as examples of the extremes of the distribution. Insert map to the bottom right shows locations of example sites. **B)** Squared factor loadings of each environmental variable for the first five dimensions. Environmental variables are described in Table 1. **C)** Histograms of the first five dimensions. **D)** Scatter plots comparing the first five dimensions. Colours match panel A. **E)** Scree plot of variance explained over all dimensions. The grey line is expected variance explained by a broken stick model.

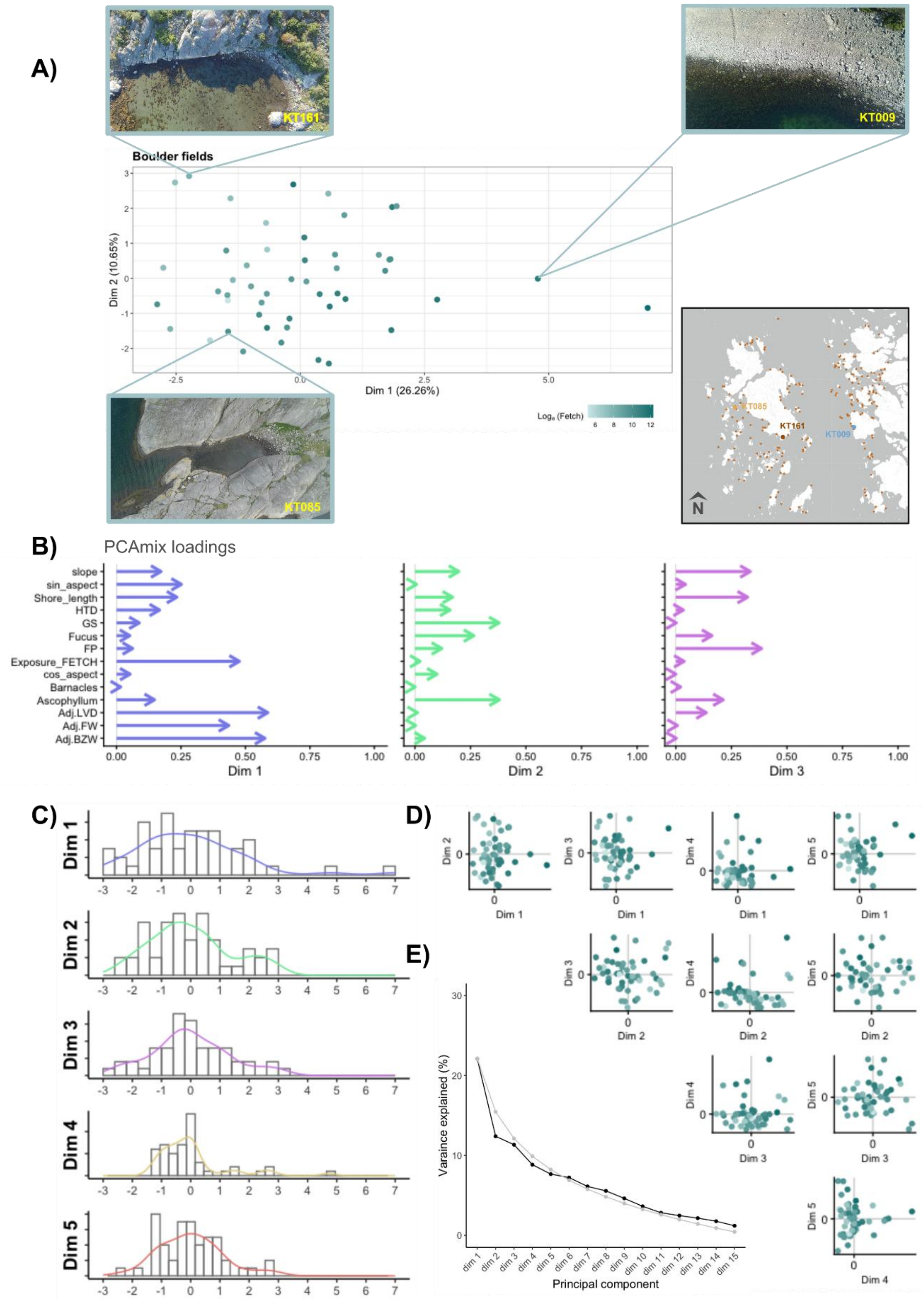


**Figure S2:** PCAmix results for all boulder shores. **A)** Scatterplot comparing PCAmix dimensions 1 and 2. Points are coloured by exposure categories observed in the field. Three drone figures are shown as examples of the extremes of the distribution. Insert map to the bottom right shows locations of example sites. **B)** Squared factor loadings of each environmental variable for the first five dimensions. **C)** Histograms of the first five dimensions. **D)** Scatter plots comparing the first five dimensions. Colours match panel A. **E)** Scree plot of variance explained over all dimensions. The grey line is expected variance explained by a broken stick model.

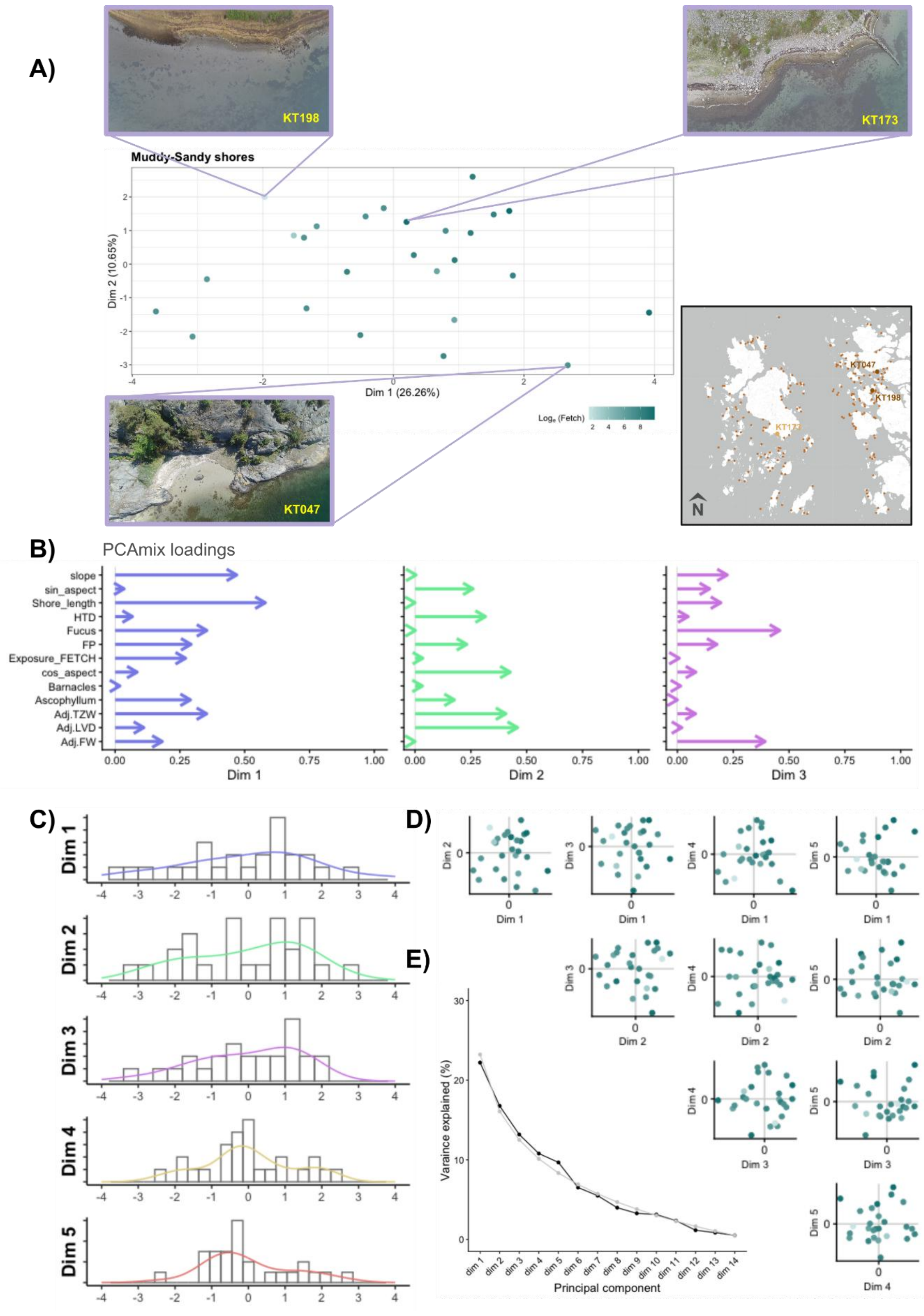


**Figure S3:** PCAmix results for all muddy and sandy shores. **A)** Scatterplot comparing PCAmix dimensions 1 and 2. Points are coloured by exposure categories observed in the field. Three drone figures are shown as examples of the extremes of the distribution. Insert map to the bottom right shows locations of example sites. **B)** Squared factor loadings of each environmental variable for the first five dimensions. **C)** Histograms of the first five dimensions. **D)** Scatter plots comparing the first five dimensions. Colours match panel A. **E)** Scree plot of variance explained over all dimensions. The grey line is expected variance explained by a broken stick model.

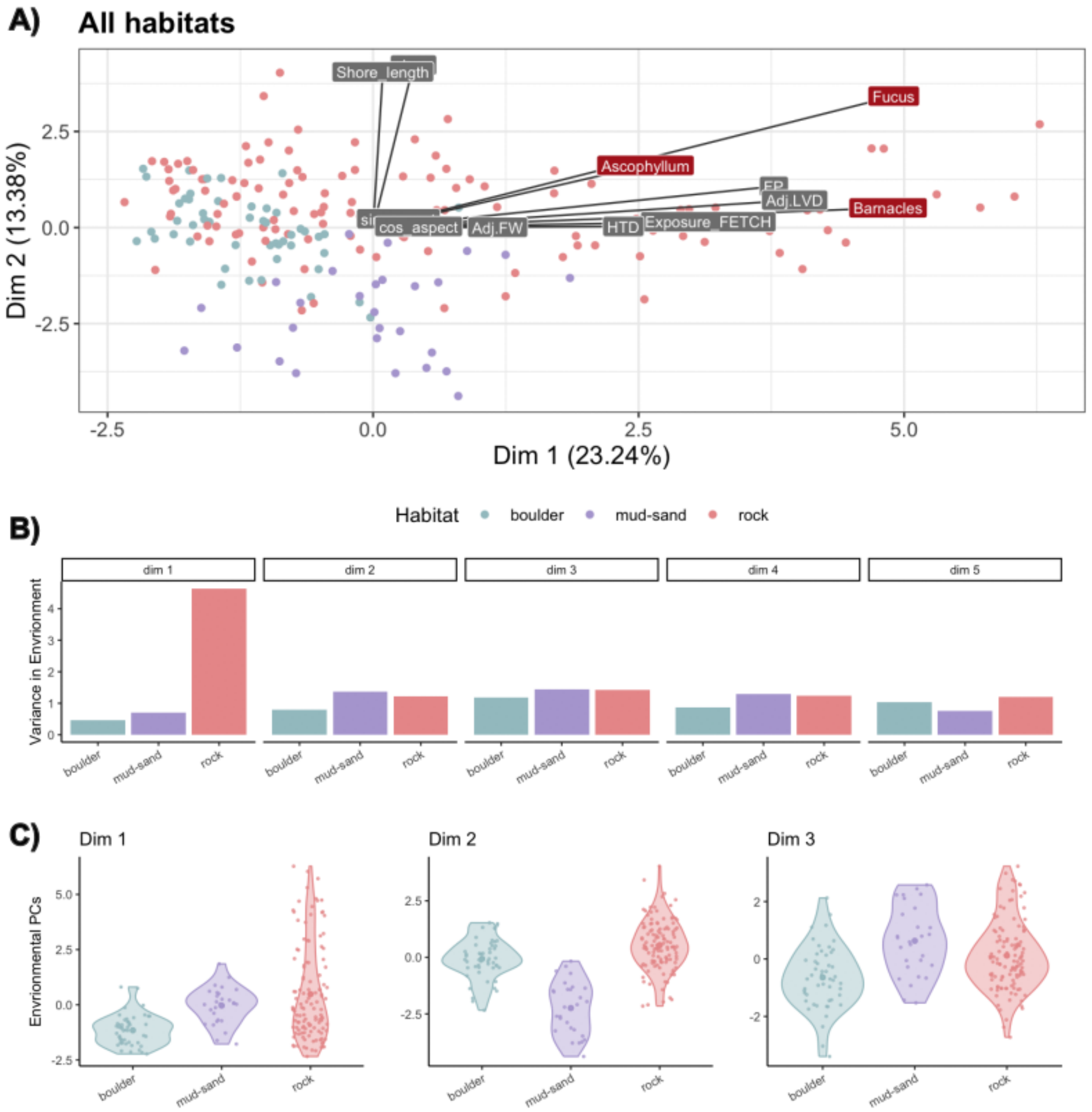


**Figure S4:** environmental variation among three habitat types. **A)** PCAmix scatterplot using only environmental measurements which were common across all habitats (see supplementary method 6). The factor loadings (eigenvectors) of each measurement are labelled above the points. Red labels indicate categorical variables. **B)** variance of environmental dimensions 1-5 within each habitat. **C)** violin plots of environmental dimensions 1-3, divided by habitats. Small points indicate individual sites. Their x-axis positions have been jittered for clarity. The larger solid dots are the mean score of each dimension.

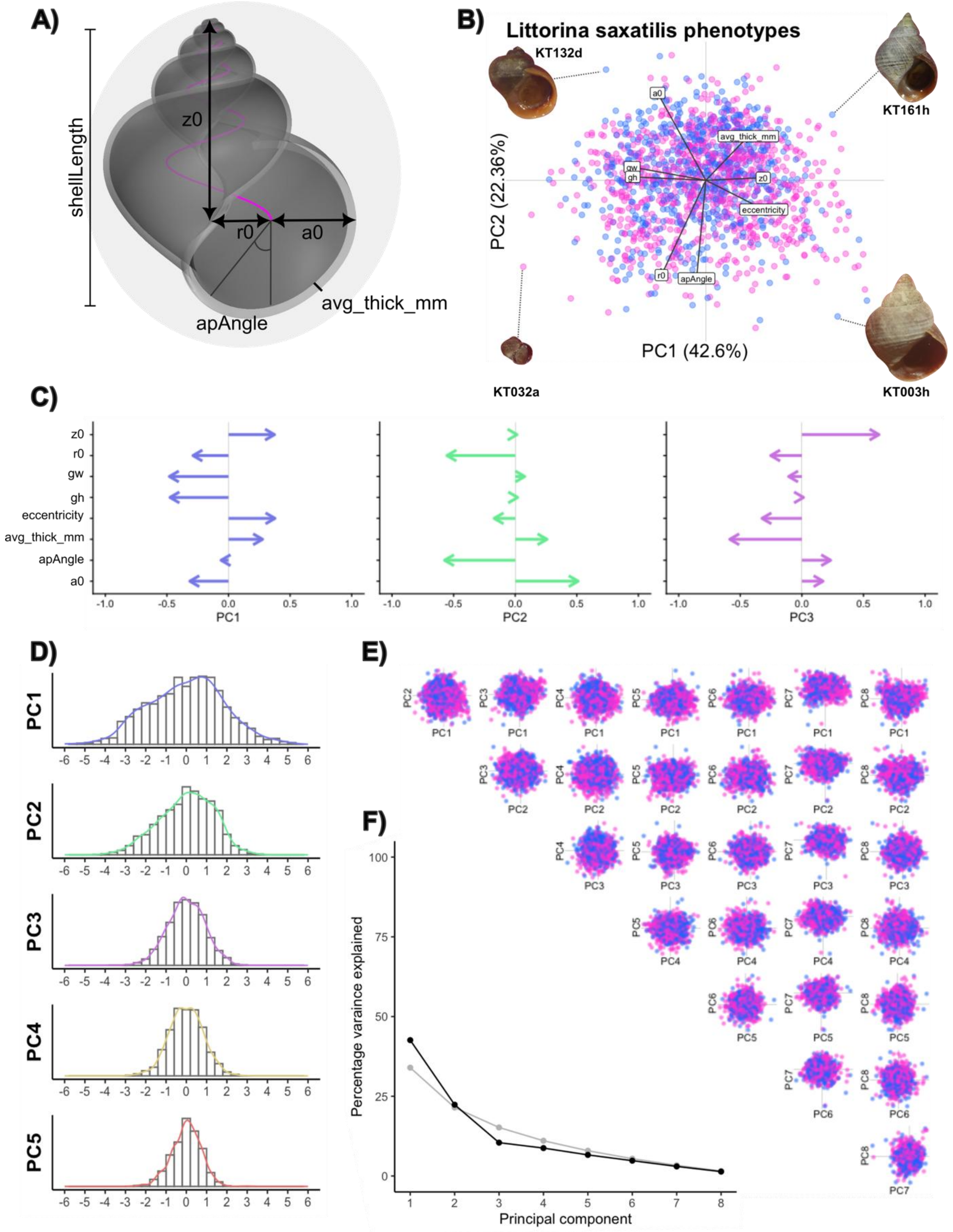


**Figure S5:** PCA results for all 1,488 snails. **A)** Diagram of a ShellShaper model indicating measurements of shells. The 'growth parameters' gw and gh are model based estimated of the spire whorls. **B)** PC1 vs. PC2 scatter plot. Points are coloured by sex (pink = female; blue = male). Labels are the factor loadings of each phenotypic variable. Four shell photos surrounding the plot. These were selected as the extreme edges of the principal components. **C)** Factor loadings over the first three principal components. Phenotypic variables are described in Table 2. **D)** Histograms of the first five principal components. **E)** PCA scatter plots comparing all eight principal components. Colours match panel B. **F)** Scree plot of variance explained over all principal components. The grey line is expected variance explained by a broken stick model.

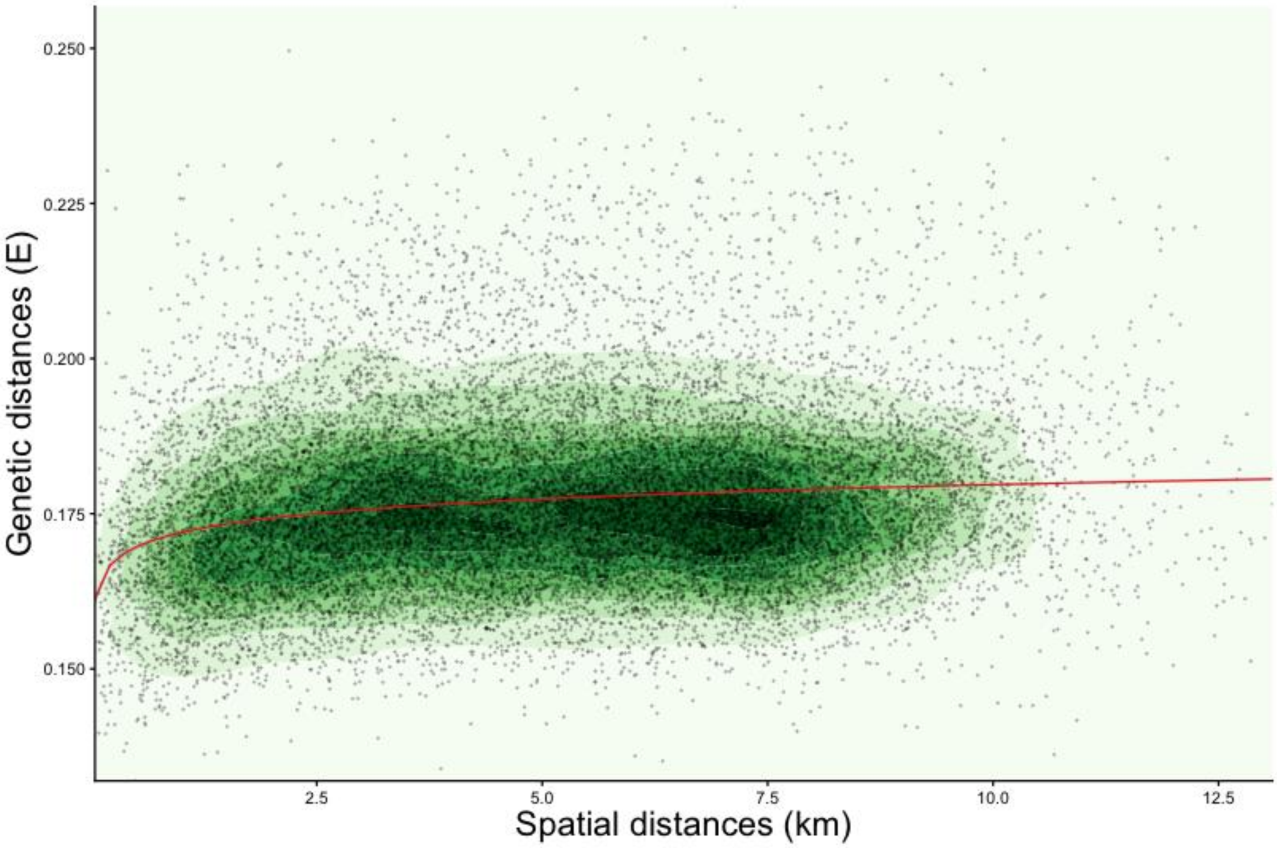


**Figure S6:** isolation by distance demonstrated by the relationship between spatial and genetic distances among sites. Points represent site pairs, and colours are the density contours estimated by *geom_denisty_2d_filled()* in the *ggplot2* R package (Wickham, 2016). Darker shades of green indicate high density. The red regression line shows the relationship between genetic distance and log-spatial distance ($\beta$ = 0.003).

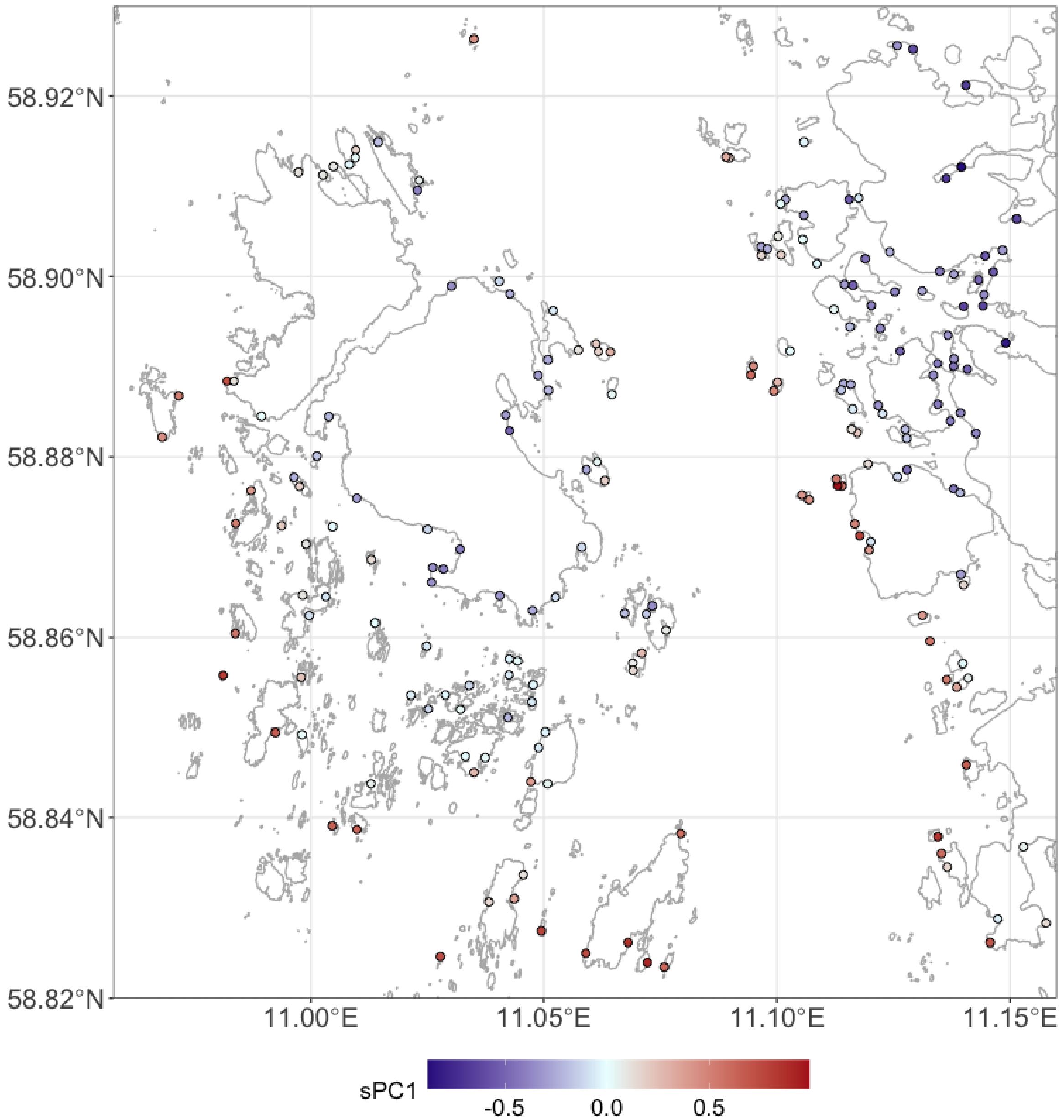


**Figure S7:** isolation by distance indicated by the first component of a spatial PCA (sPC1) calculated with the *adgenet* R package (Jombart, et al., 2008; 2011). Similar colours show genetically similar sites. The basemap was taken from the coastline vector from the Swedish Land Survey (see Fig. 1 for details).

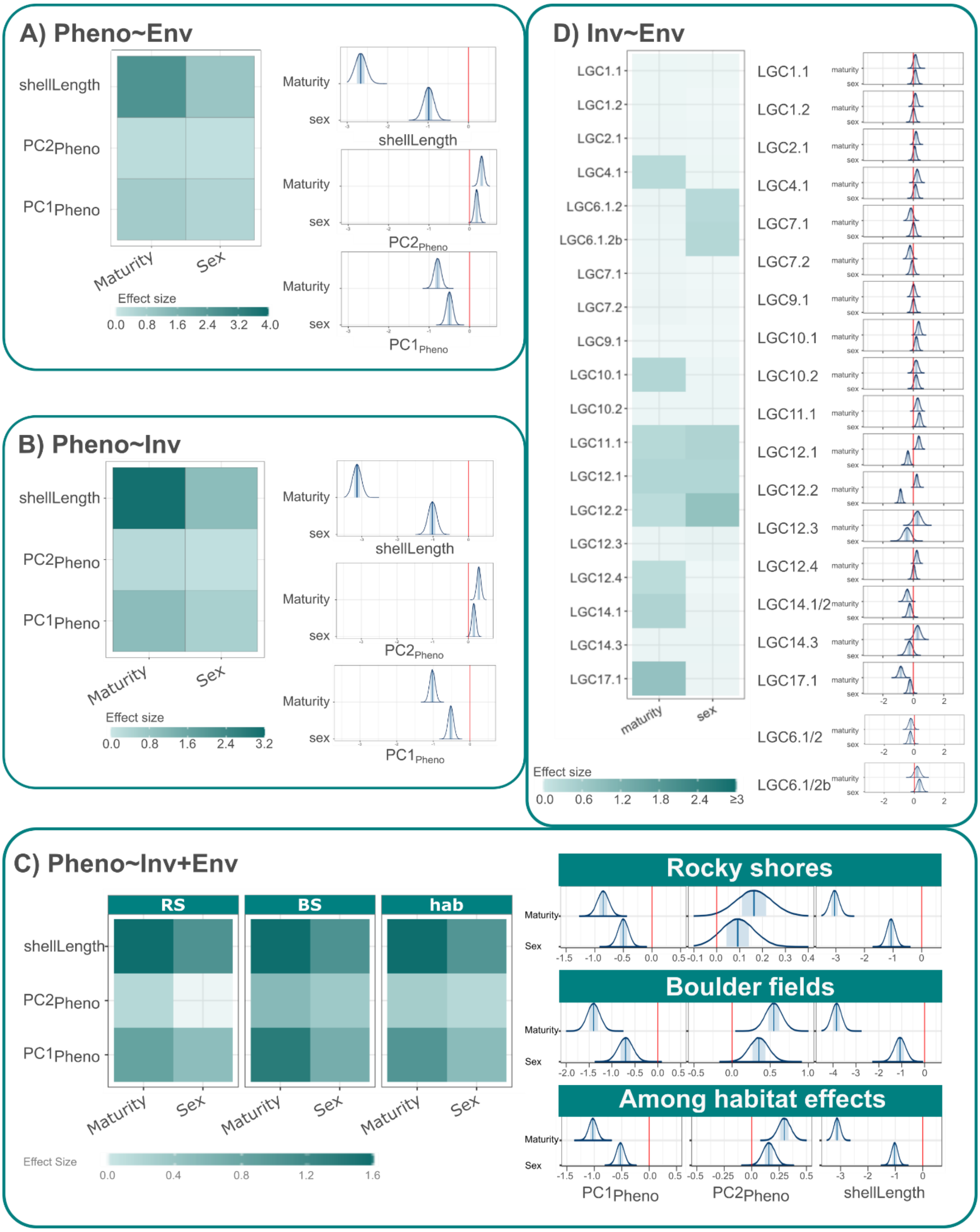


**Figure S8:** summary of the effects of maturity and sex for **A) Pheno~Env**, **B) Pheno~Inv**, **C) Pheno~Inv+Env**, and **D) Inv~Env** Bayesian regression models. Tile plots summarise the effect size. Darker tiles are stronger effects. Filled tiles indicate informative effects, while crossed tiles are uninformative. Posterior distributions are shown to the right of each tile plot. Vertical blue lines indicate the median and the blue box indicates the 50% interval. Only the results of among habitat models, which use all samples, are considered for maturity and sex effects in **Pheno~Env** models.

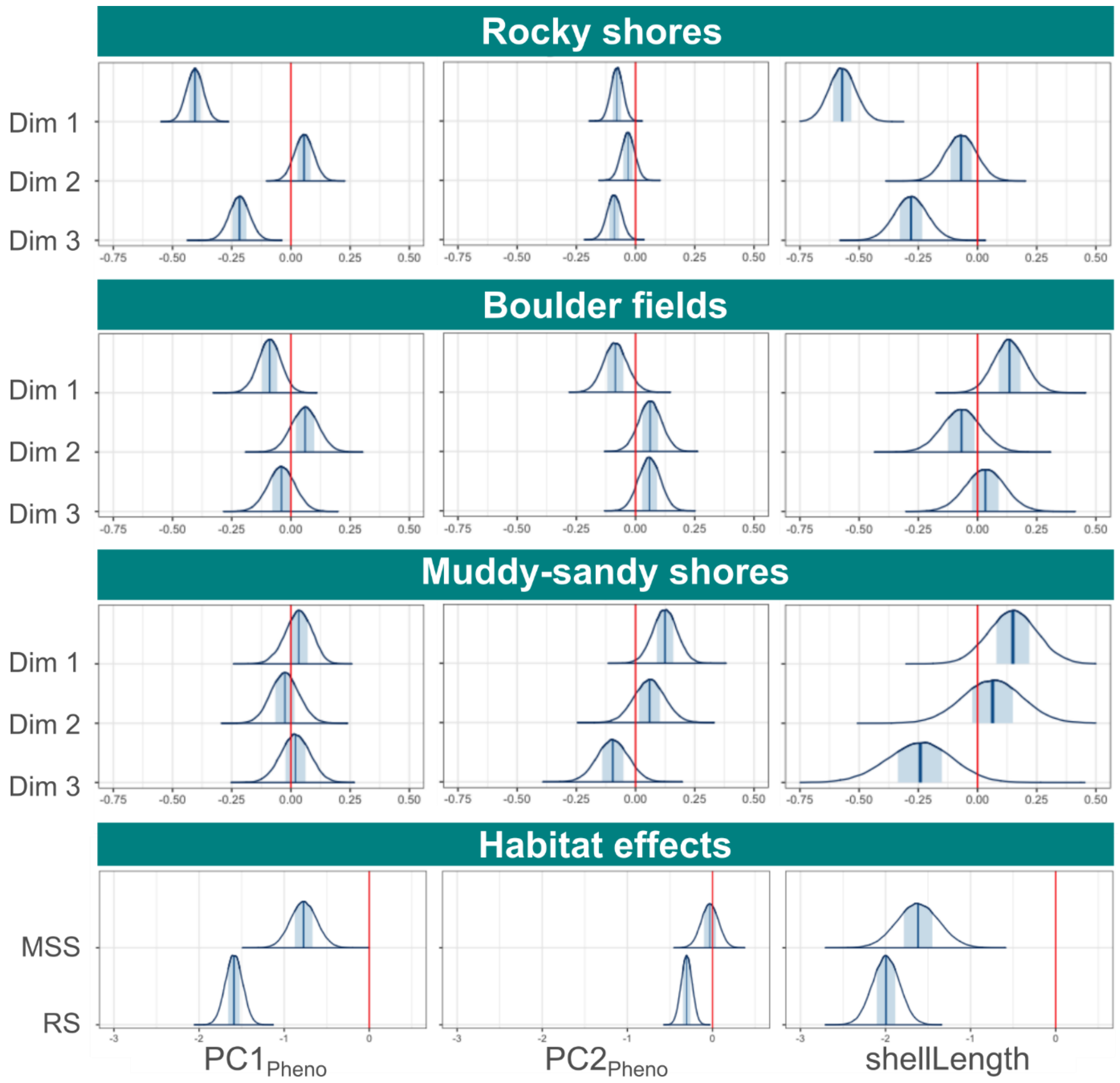


**Figure S9:** posterior distributions for **Pheno~Env** Bayesian regression models. 'Dim' are the environmental PCAmix dimensions. Vertical blue lines indicate the median and the blue box indicates the 50% interval. Habitat effects show the difference between boulder shores and either muddy-sandy shores (**MSS)** or rocky shores (**RS**). Tile plot summary for these fits is in the main text (Fig. 5A).

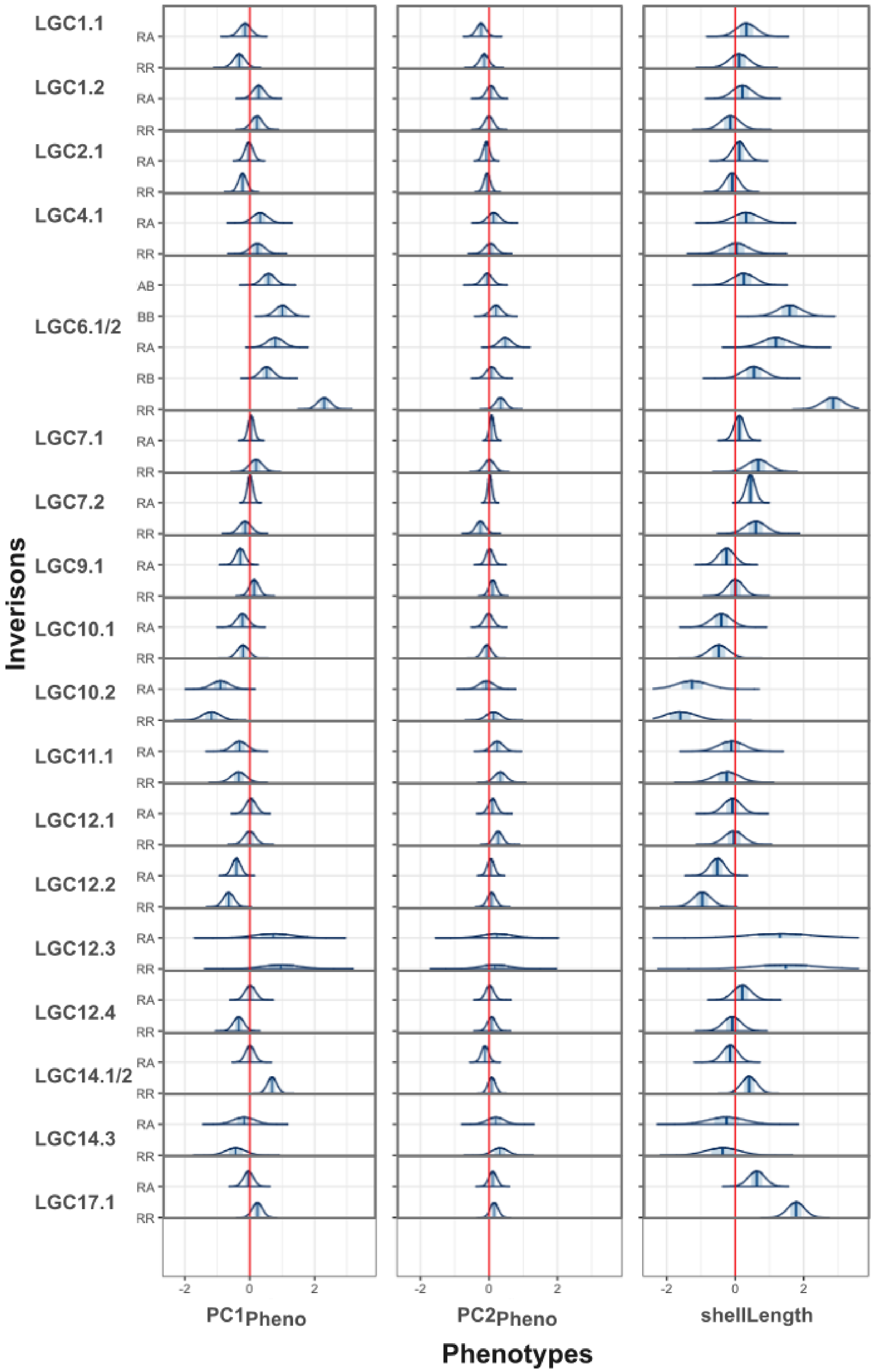


**Figure S10:** posterior distributions for **Pheno~Inv** Bayesian regression models. Vertical blue lines indicate the median and the blue box indicates the 50% interval. Tile plot summary for these fits is in the main text (Fig. 5C). Posterior estimates are all made as the difference from the mean phenotypic score of the 'AA' genotype.

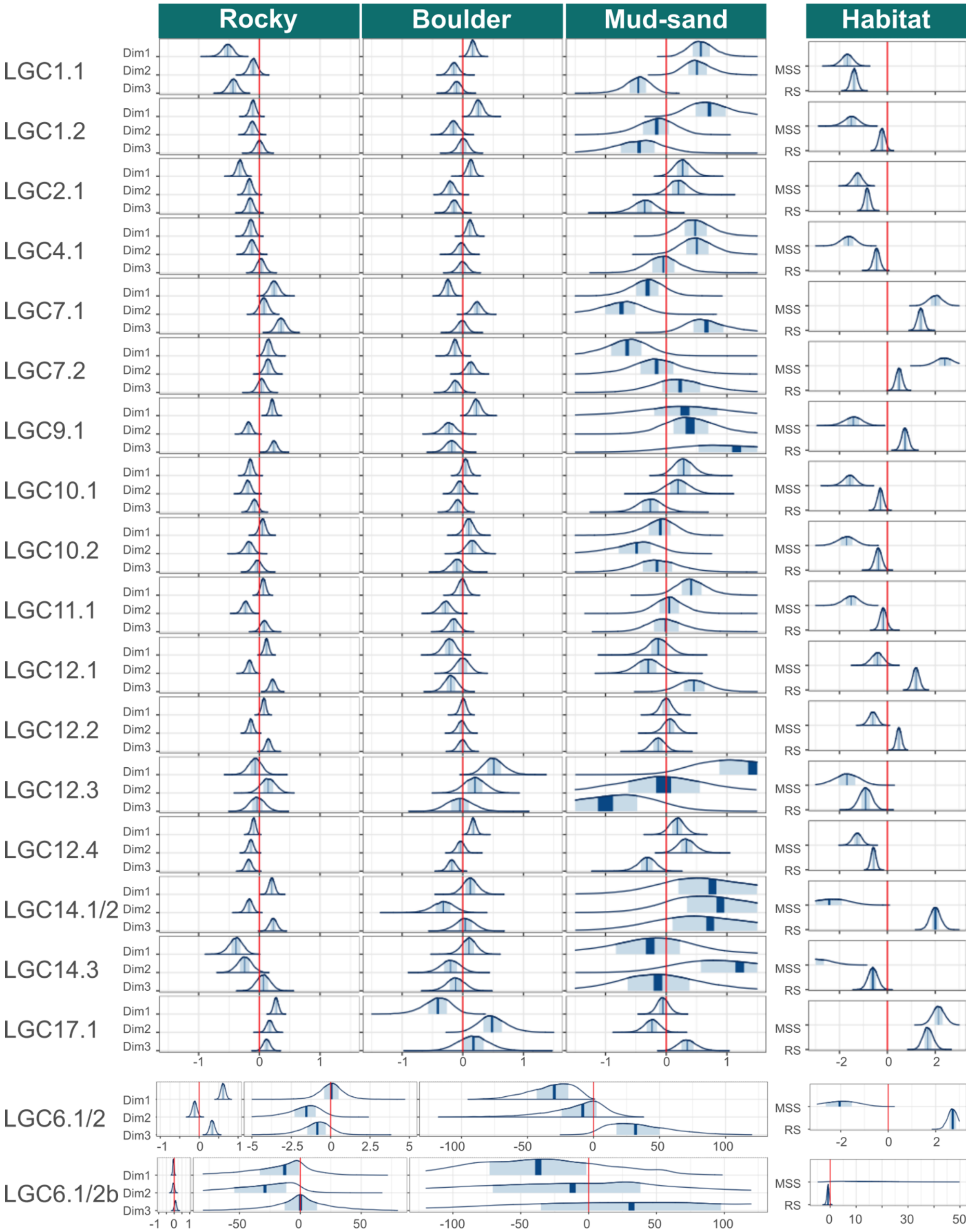


**Figure S11:** posterior distributionss for **Inv~Env** Bayesian regression models. Vertical blue lines indicate the median and the blue box indicates the 50% interval. 'Dim' are the environmental PCAmix dimensions. 'Habitat' show the difference between boulder shores and either muddy-sandy shores (**MSS)** or rocky shores (**RS**). Panels for LGC6.1/2 have asymmetric x-axes to emphasise the different levels of uncertainty among models. Tile plot summary for these fits is in the main text (Fig. 5E).

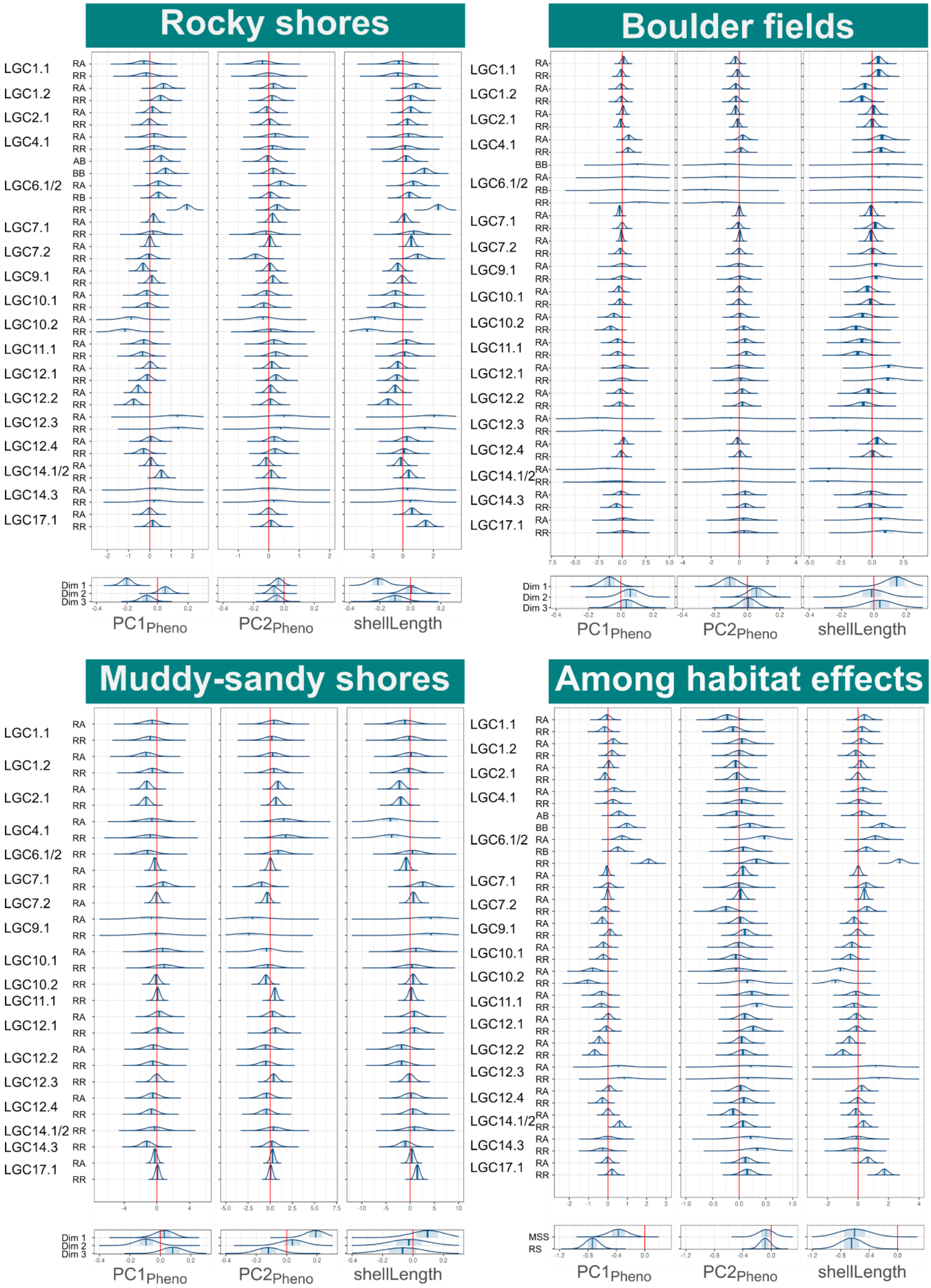


**Figure S12:** Posterior distributions for **Pheno~Inv+Env** Bayesian regression models. Vertical blue lines indicate the median and the blue box indicates the 50% interval. Some distribution edges have been pruned to provide a better resolution to other distributions on a common axis. Inversion and environmental effects are in separate panels. Columns show the different habitat classes. Tile plot summary for these fits is in the main text (Fig. 6).

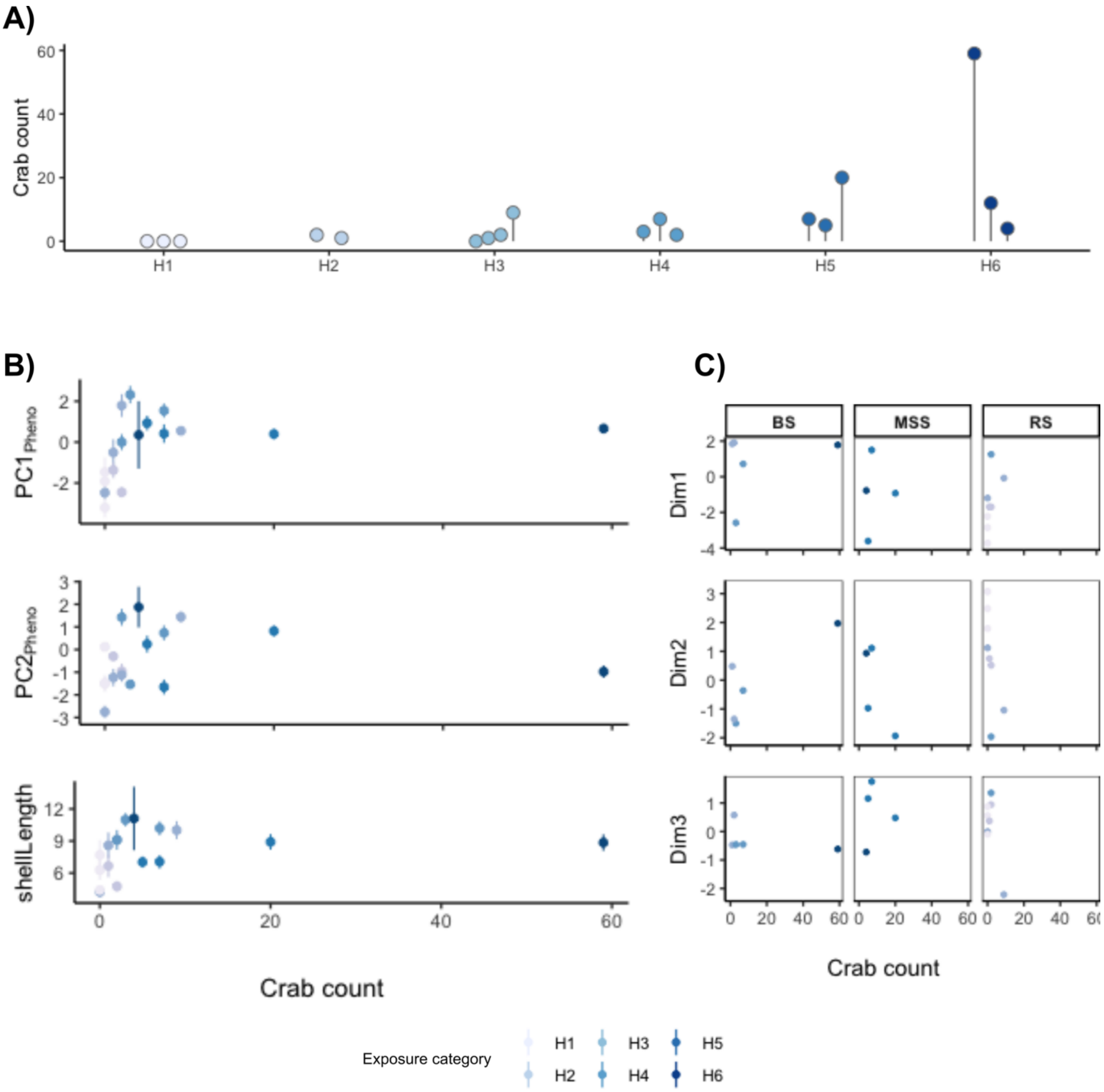


**Figure S13:** Crab density estimates from a pilot study at 18 sites (Table S3). **A)** Count of crabs, summed over six observations, per exposure category. H1 are extremely exposed and H6 are highly sheltered sites. Each point represents a separate sampling site. Counts are the total of six trials at each site. **B)** Effects of crab density on phenotypic PCs. Points are the site average PC score, with error bars indicating the standard error. **C)** Effects of crab density on environmental variables. Points are the Dim score for each site. Dim was calculated separately for each habitat class: **BS** = boulder shores, **MSS** = muddy-sandy shores, and **RS** = rocky shores.

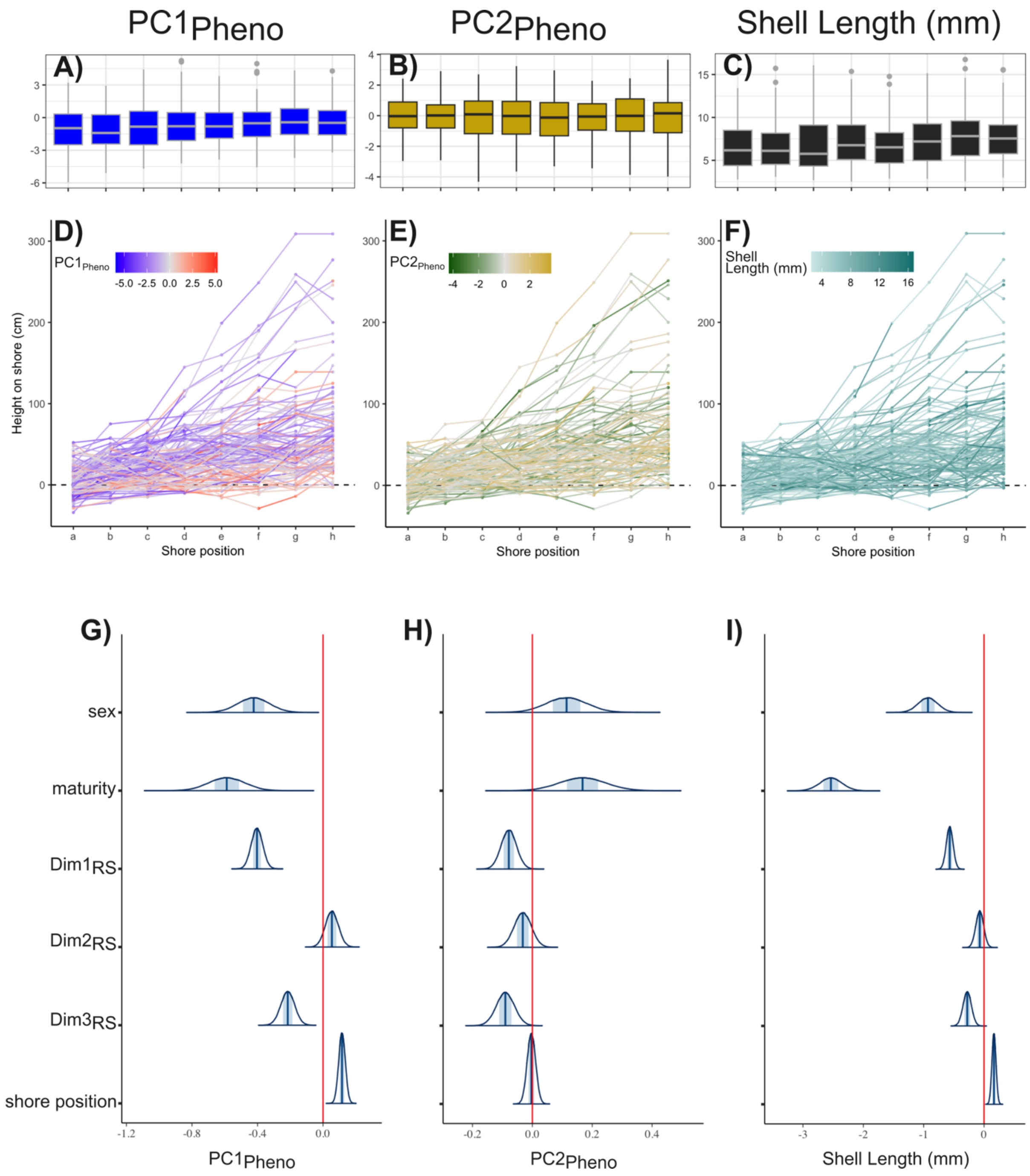


**Figure S14:** Effects of adding shore position of each snail to rocky shore **Pheno~Env** models. Shore positions are categories representing different points along the transect, with 'a' being the lowest and 'h' highest point shore. **A-C**) boxplots of phenotypic scores for each shore position. Median is shown as the central tendency. **D-F)** height on shore (cm) each snail was sampled from, grouped by shore position. Heights were adjusted by average sea level. Points are coloured by phenotypic score per snail. Lines connect snails collected from the same site. **G-I)** posterior distributions for individual-level (i.e., fixed) parameter estimates.

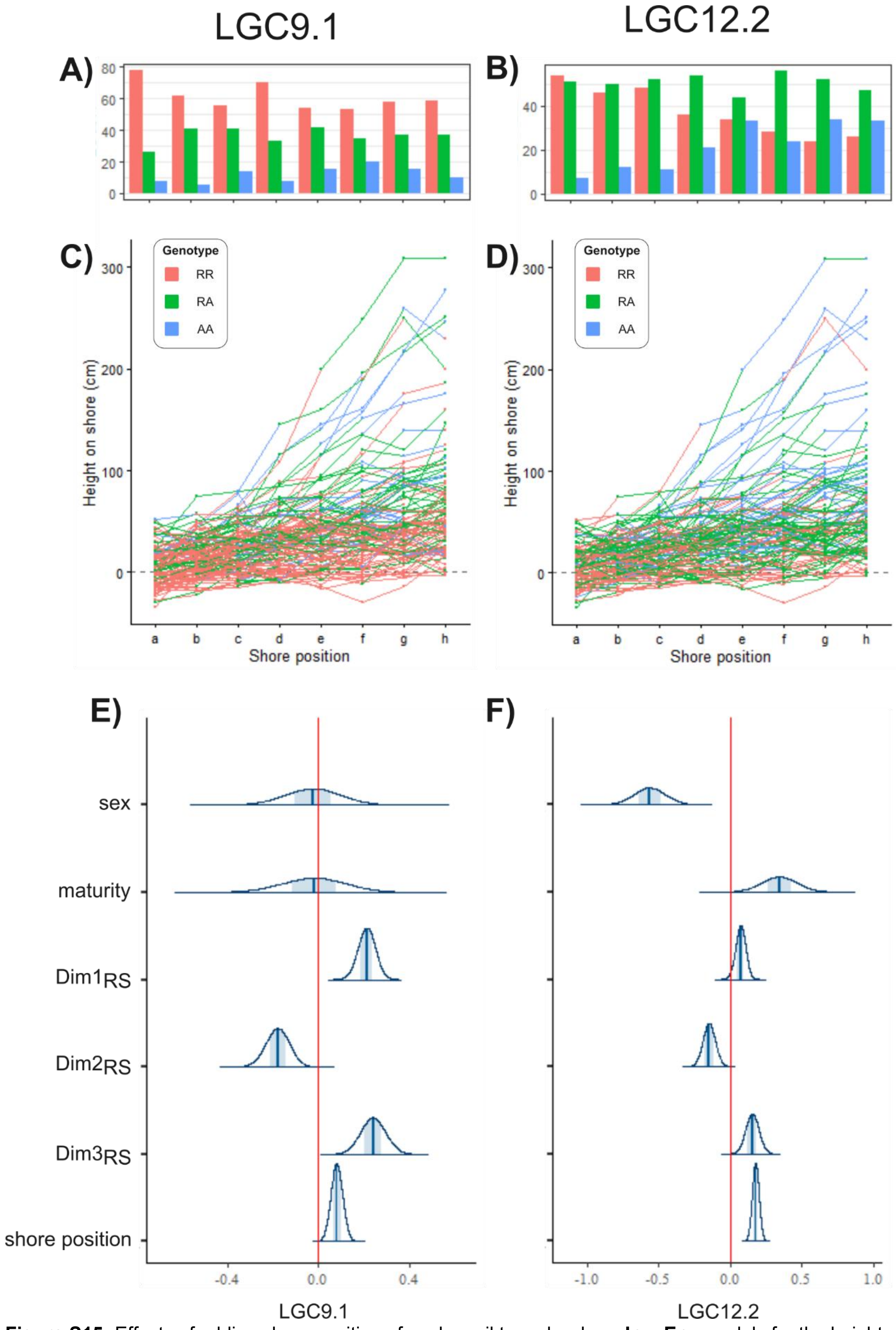


**Figure S15:** Effects of adding shore position of each snail to rocky shore **Inv~Env** models for the height associated inversions; LGC9.1 and LGC12.2. Shore positions are categories representing different points along the transect, with ‘a’ being the lowest and ‘h’ highest point shore. **A-B**) barplot of genotypes for each shore position. **C-D)** height on shore (cm) each snail was sampled from, grouped by shore position. Heights were adjusted by average sea level. Points are coloured by each snail’s genotype. Lines connect snails collected from the same site. **E-F)** posterior distributions for individual-level (i.e., fixed) parameter estimates.

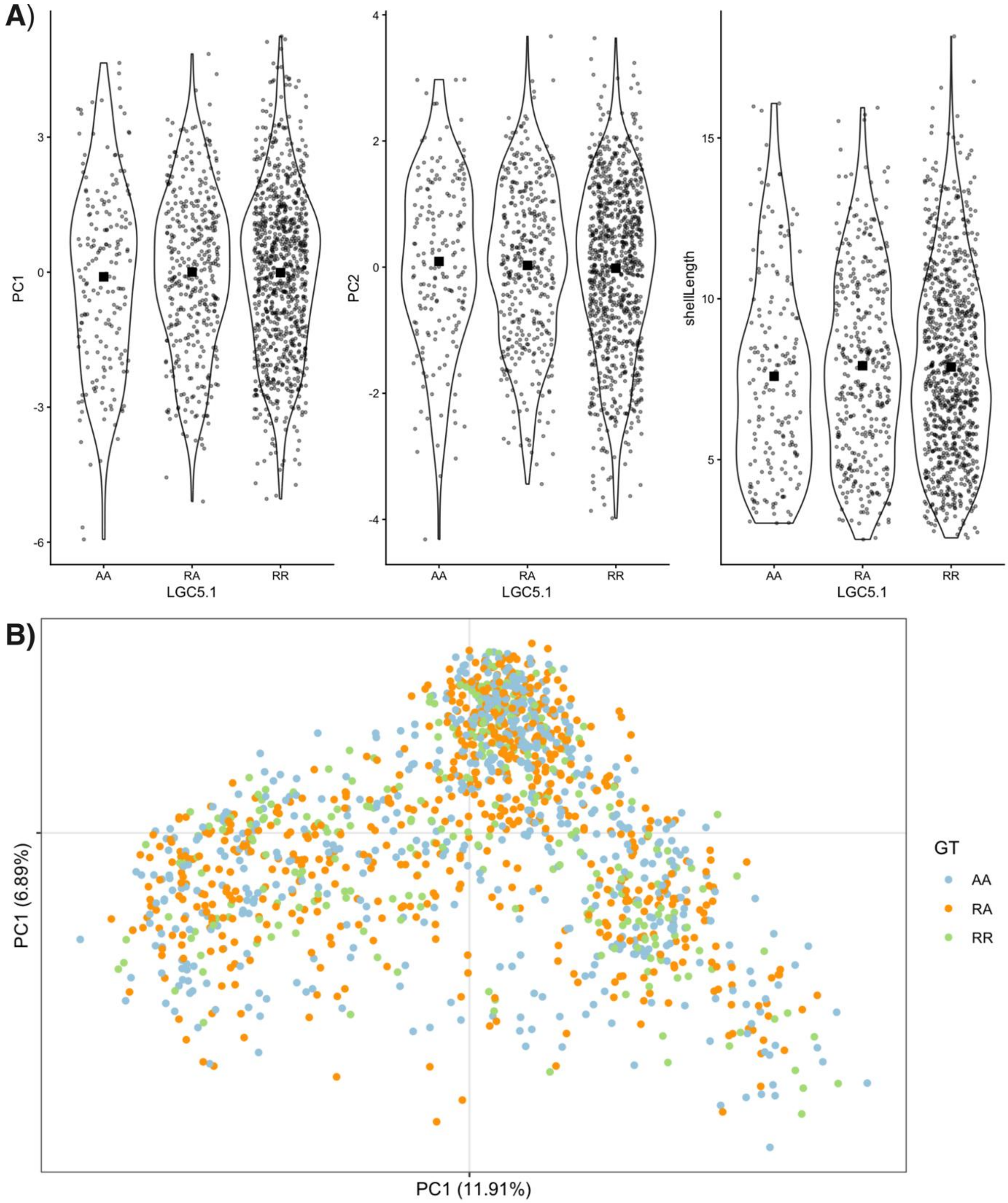


**Figure X:** influence of LGC5.1 on phenotypic variables and genetic background. **A)** violin plots of phenotypic variables split by LGC5.1 genotype. Small points indicate individual samples. Solid black squares indicate the means. **B)** PCA scatterplot for the first two axes genetic background SNPs. Points are coloured by LGC5.1 genotypes.
